# DeepVariant calling provides insights into race diversity and its implication for sorghum breeding

**DOI:** 10.1101/2022.09.06.505536

**Authors:** Pradeep Ruperao, Prasad Gandham, Damaris A Odeny, Sivasubramani Selvanayagam, Nepolean Thirunavukkarasu, Roma R Das, Manasa Srikanda, Harish Gandhi, Ephrem Habyarimana, Eric Manyasa, Baloua Nebie, Santosh P Deshpande, Abhishek Rathore

## Abstract

Due to evolutionary divergence, sorghum race populations exhibit vast genetic and morphological variations. A *k-mer*-based sorghum race sequence comparison identified the conserved *k-mer*s of all sorghum race accessions and the race-specific genetic signatures identified the gene variability in 10,321 genes (PAVs). To understand the sorghum race structure, diversity and domestication, deep learning-based variant calling approach was employed in a set of genotypic data derived from a diverse panel of 272 sorghum accessions. The data resulted in 1.7 million high-quality genome-wide SNPs and identified selective signature (both positive and negative) regions through a genome-wide scan with different (iHS and XP-EHH) statistical methods. We discovered 2,370 genes associated with selection signatures including 179 selective sweep regions distributed over 10 chromosomes. Localization of these regions undergoing selective pressure with previously reported QTLs and genes revealed that the signatures of selection could be related to the domestication of important agronomic traits such as biomass and plant height. The developed *k-mer* signatures will be useful in the future to identify the sorghum race and SNP markers assist in plant breeding programs.

## Introduction

The process of domestication and natural selection leads to an increased frequency of favourable alleles and subsequently results in complete fixation at target genomic loci (Smýkal *et al*., 2018). Although the selection process targets advantageous alleles, it also inadvertently results in an increase in the frequency of alleles at the neutral loci that are in linkage disequilibrium, a phenomenon referred to as selective sweep (Stephan *et al*., 1992). Selective sweep has the potential of enhancing the fitness of an individual at the expense of the overall genetic diversity of a population at the respective loci. As a result, modern cultivars are derived from a small fraction of genetically related varieties (Mccouch *et al*., 2013) in spite of the existence of the vast genetic diversity of global plant germplasm. A better understanding and exploitation of existing natural variation in each crop is a key aspect of meeting the increasing food demand in the coming decades.

Sorghum *[Sorghum bicolor* (L.) Moench] is an important cereal crop grown and consumed by a majority of the global population. The earliest record of sorghum seeds that were found at Nabta Playa (Egyptian-Sudanese border) indicates early domestication (Wendorf *et al*., 1992). The subsequent migration and adaptation of sorghum across Africa and Asia led to the evolution of morphological and geographically diverse groups, classified into major races (Harlan and Wet, 1972; Harlan and Stemler, 2012). The more recent phenotype and genotype-based classifications also support the sorghum race classification within the global diversity panel (Brown *et al*., 2011). However, inter-racial diversity has not been fully understood in sorghum in a way that deprives our understanding of the racial structure and potential heterotic gains. Development of such knowledge would improve overall genomic predictions in sorghum as has been done in other cereal crops (Norman *et al*., 2018) for the best use of the genome in crop improvement programs.

The extent of genetic diversity is measured by the number of nucleotide variations across individuals and species (Deu *et al*., 2006; Kebbede, 2020). Such variations range from single nucleotides to large-scale structural variations. However, most studies in the past have only used single nucleotide variations (Afolayan *et al*., 2019; Enyew *et al*., 2022) ignoring other structural variations such as insertion-deletions (indels) and presence-absence variations (PAV) (Saxena *et al*., 2014). PAVs are present in some individuals but absent in others, making them perfect for detecting major differences among multiple genomes. Pangenomes, therefore, can help obtain the complete genomic variants of a species (Hurgobin and Edwards, 2017) since they hold the exhaustive number of genes of a given species. The availability of sorghum pangenomes (Ruperao *et al*., 2021a; Tao *et al*., 2021) makes it possible to carry out a more extensive genetic variation analysis across the different races.

Despite emerging advancements in sequencing technologies, calling accurate genetic variants from sequencing errors remains challenging. Because a majority of the genome assembly tools are based on the *de Bruijn* graphs (Bankevich *et al*., 2012; Peng *et al*., 2012; Simpson *et al*., 2009; Zerbino and Birney, 2008), in which the sub-sequence of *k-mer*s (substring of length *k*) are used to construct the graph and output the paths as contigs (without branching). The resulting contigs can therefore be biased and fragmented as a result of sequencing errors, especially in highly repetitive genomes, leading to low confidence in variant calling. Alternative alignment-free methods of variant detection have been developed using both *k-mer* frequencies and information theory (Song *et al*., 2014; Pajuste *et al*. 2017; Zielezinski *et al*. 2017; Audano *et al*. 2018). These alignment-free methods have been applied in several studies including phylogeny estimation (Haubold, 2014), identification of mutations between strains (Nordström *et al*., 2013) and association mapping (Sheppard *et al*., 2013).

More recently, deep learning methods have been introduced as a machine learning technique applicable to a range of fields including genomics. Deep learning models can be trained without prior knowledge of genomics and next-generation sequencing (NGS) data to accurately call genetic variants (Telenti *et al*., 2018). Learning a deep convolutional neural network-based statistical relationship between the aligned reads, genotypes calling approach has been implemented in DeepVariant programs (Poplin *et al*., 2018a). The DeepVariant approach is reported to outperform the existing variant calling tools (Poplin *et al*., 2018b).

The objective of our study was to use deep learning (DeepVariant method) to better understand genetic variation, domestication events and selection signatures across known sorghum races. We used existing whole-genome sequence data to quantify genome-wide positive and negative selection regions to enhance our understanding of the genome function and the frequency of genetic variations. In addition, we determined the putative signals of selection in sorghum have resulted from true selective events or population bottlenecks.

## Results

### Access to raw data

We obtained publicly accessible raw Illumina sequence data from three previous studies as shown in Table 1 and Supplementary Table 1. The sorghum accessions having minimum 5x coverage of whole-genome sequence data were used for the analysis, resulting in a total of 272 sorghum accessions.

**Table 1.** A summary of publicly available data used in the current analysis.

| Reference | # Genotypes | Average coverage |
| --- | --- | --- |
| Valluru *et al.* (2019) | 196 | 13x |
| Jensen *et al.* (2019) | 70 | 7x |
| Yan *et al.* (2018) | 6 | 28x |
| **Total** | **272** | **16x** |

### DeepVariant calling and annotation

The whole genome sequence (WGS) data were mapped (Supplementary Table 2, Supplementary Figure 1) to the sorghum pangenome (Ruperao *et al*., 2021), and a total of 1.7 million high-quality SNPs, and 470,375 InDels (154,900 insertions, 278,951 deletions and 36,524 mixed variants) were called using the deepVariant method. Homozygous SNPs were predominant (88.3%) over heterozygous SNPs (11.6%) (Supplementary Table 3). The overall density of SNPs was 2.5 SNP/kb, whereas the indels density was 0.6/kb. The maximum (209,429) and minimum (147,952) SNPs were reported on chromosome 2 (0.3/kbp) and chromosome 9 (0.4/kbp), respectively (Supplementary Table 4) (Figure 1), while the maximum (70,722) and minimum (34,530) number of indels were reported on chromosome 1 (0.8%) and chromosome 8 (0.5%) respectively. Most of the insertions (98%) and deletions (93%) were less than 10bp in length (Figure 2.a).

**Figure 1:**
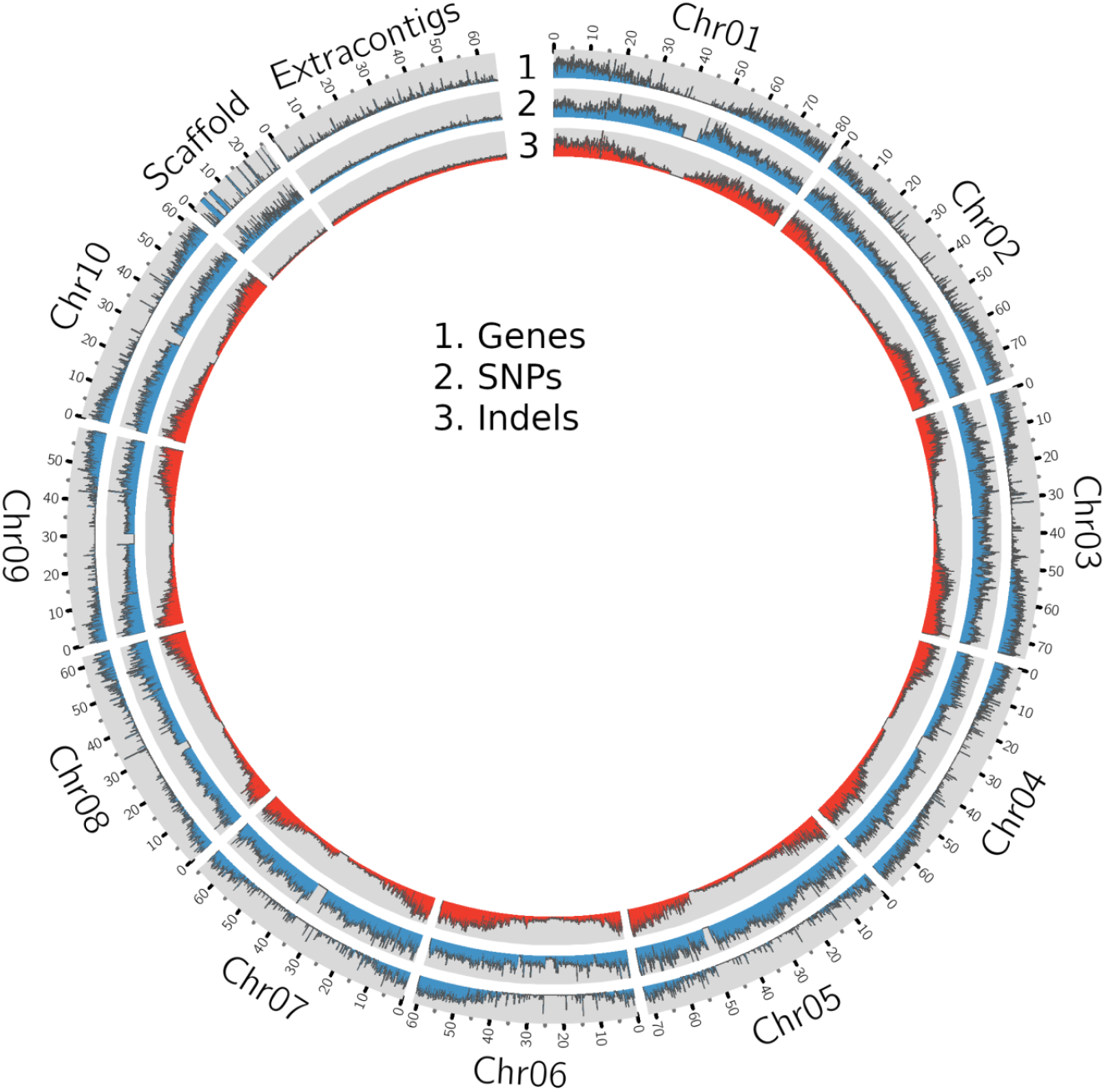
A Circos plot showing the density of genes, SNPs and InDels in the sorghum pangenome.

**Figure 2:**
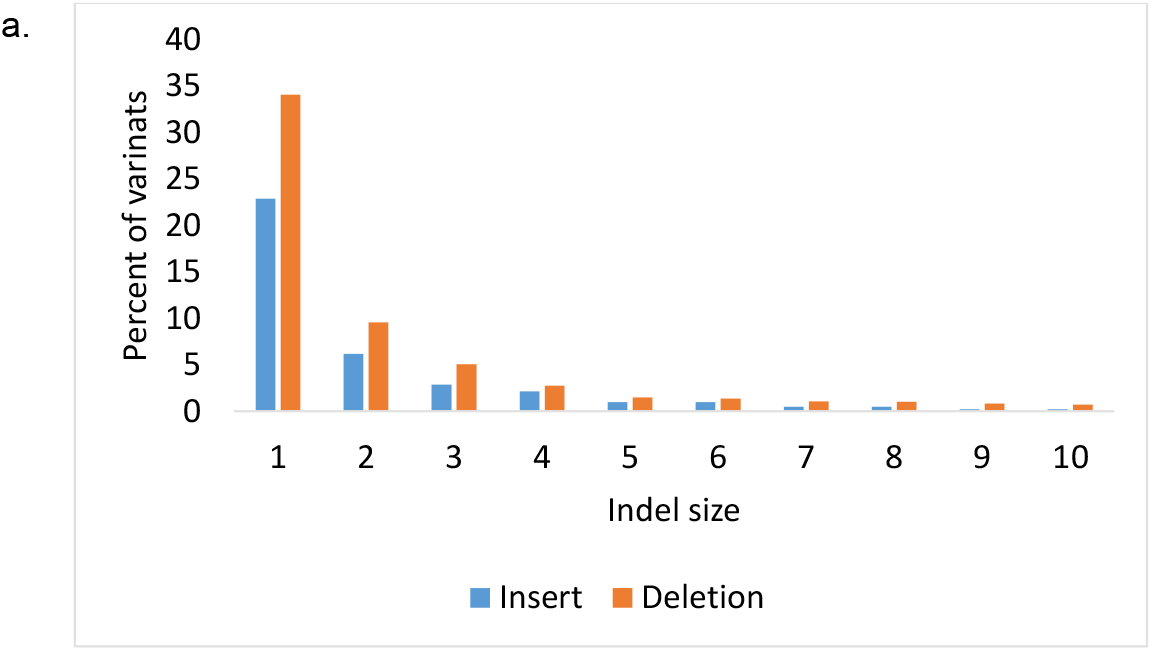
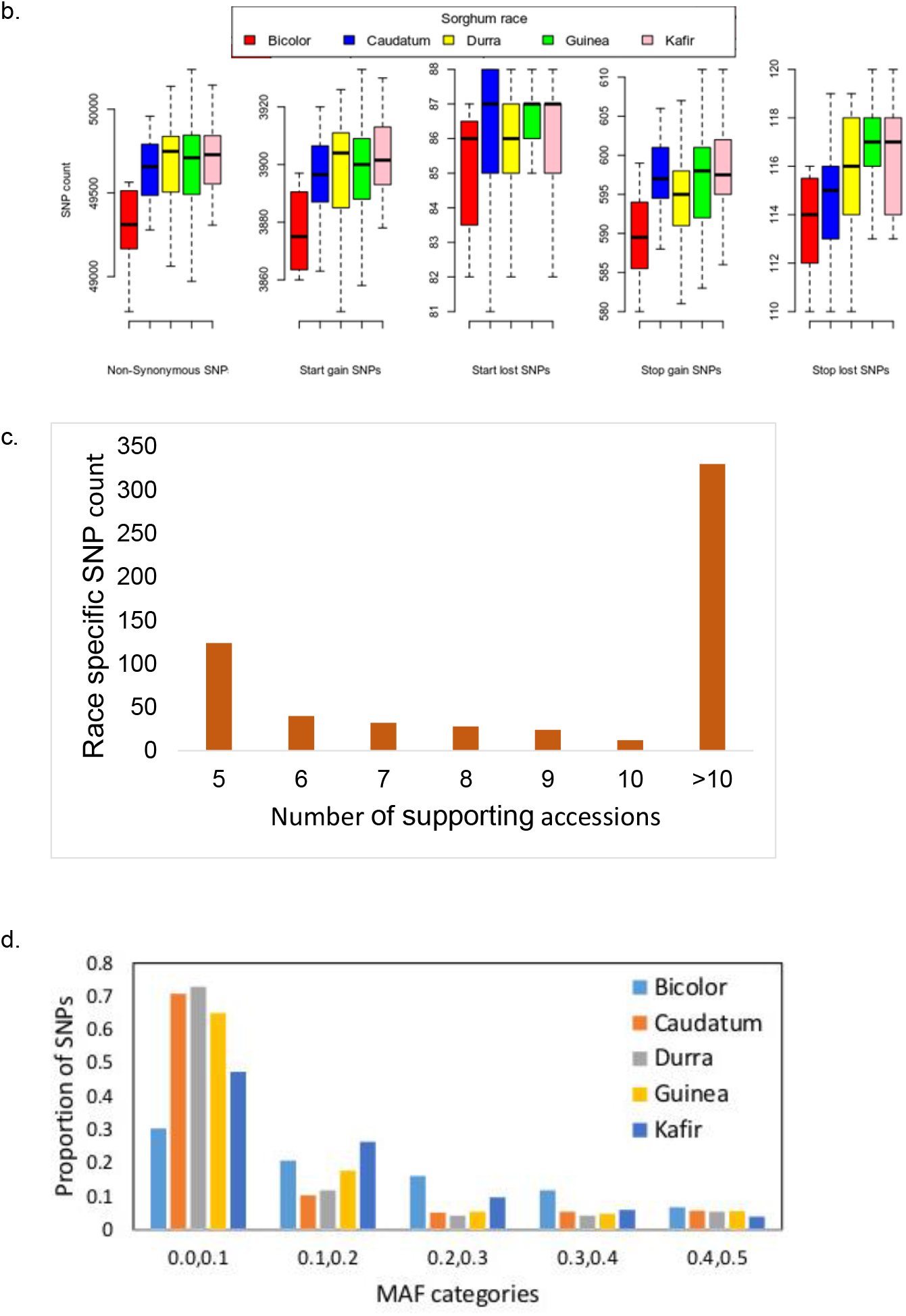
Genomic landscape of sorghum. a) Indel length distribution b) SNP annotations c) race-specific SNP calls with a number of supporting sorghum accessions d) The distribution of minor allele frequencies for all sorghum races. SNPs were binned into five categories on the MAF.

The SNP annotation reported 11% of SNPs from of which 51,891 were synonymous and 53,159 non-synonymous, resulting in a non-synonymous-to-synonymous substitution ka/ks ratio of 1.02 (Supplementary Tables 5 and 6), consistent with the previous study by Mace *et al*. (2013). Sorghum accessions NSL54318 (50,238 non-synonymous; 3,933 start gain; 88 start lost and 611 stop gain SNPs) and PI660645 (46520 non-synonymous; 3700 start gained and 103 stop lost) harboured the maximum and minimum effect SNPs (Supplementary Table 5). There were more transitions (C/T and A/G) than transversions (A/T, A/C, T/G and C/G) with a transition/transversion ratio ranging from 1.912 (NSL50716, IS30508) to 1.983 (PI329719). The overall tr/tv ratio was 1.960 (Supplementary Table 7).

SNPs with large effects were the least common (1,362; 0.04%) compared to SNPs with low (63,298; 1.9%), moderate (53,159; 1.6%) and modifying SNPs (96%). A total of 89.3% (1,595,340) of the SNPs were conserved across five sorghum race accessions while the remaining 10.6% (190,321) were variably detected in at least one sorghum race. Among the SNPs in the sorghum race accessions, 0.03% (590) were race-specific, the majority (60.6%; 358) of which were reported in durra and the least in the bicolor race (6.1%; 36) (Supplementary Table 8) (Figure 2.b). Most of the race-specific SNPs (57.9%) were highly confident with support from more than 10 accessions. Only 21% of the race-specific SNPs were supported by less than 5 accessions (Figure 2. c).

Sorghum races caudatum, durra, guinea and kafir had the highest proportion of SNPs with the low MAF category (0.0,0.1) compared to bicolor. Kafir had the highest proportion of SNPs with MAF category (0.1, 0.2) while bicolor race reported the highest proportion of SNPs with MAF greater than 0.2, which is expected for a race with a long history of cultivation (Figure 2.d)

### Genetic and nucleotide diversity

The SNP-based Neighbour-Joining (NJ) dendrogram of the 272 genotypes grouped them largely according to race genetic relatedness. Four major clusters were observed with a number of subgroups. The phylogenetic tree contained a distinct cluster of 63 guinea race accessions (nodes in blue colour) mixed with a few other race individuals, such as durra (PI221662, PI248317, PI267653 and PI148084) (nodes in brown colour), kafir (PI660555 and NSL365694) (nodes in pink colour), bicolor race (IS12697) (nodes in red colour). The other sorghum race clusters were split with non-corresponding sorghum race accessions. For example, durra has 91 accessions split into two clusters with caudatum and kafir accessions. The bicolor accessions were placed mostly in durra and guinea clusters. Among the bicolor accessions, the China origin accessions were grouped distinctly in the durra cluster compared to other bicolor accessions.

The evaluation of nucleotide diversity across all 272 accessions showed that sorghum had low diversity (0.0000483715) compared to wheat (*π_A_*=0.0017, *π_B_* = 0.0025 and *π_D_* = 0.0002) (Zhou *et al*., 2020), maize (*π* = 0.014) (Tenaillon *et al*., 2001) and rice *π* = 0.0024 (Huang *et al*., 2010) (Figure 3) (Supplementary Figure 3). The diversity varies on the population size and the level of diversity of the accessions used in such a population. However, such low diversity was also reported in an earlier study (Sapkota *et al*., 2020). We observed significant differences (*P* < 0.05) in nucleotide diversity between three sorghum races (caudatum, durra and guinea) that were represented with more than 50 genotypes. The durra had the highest nucleotide diversity while caudatum showed the lowest (*π_C_* = 0.0000419, *π_G_* = 0.0000631 and *π_D_* = 0.0000637). The distribution of nucleotide diversity on the sorghum race genome was in the order of *π_D_* > *π_G_* > *π_C_*.

**Figure 3.**
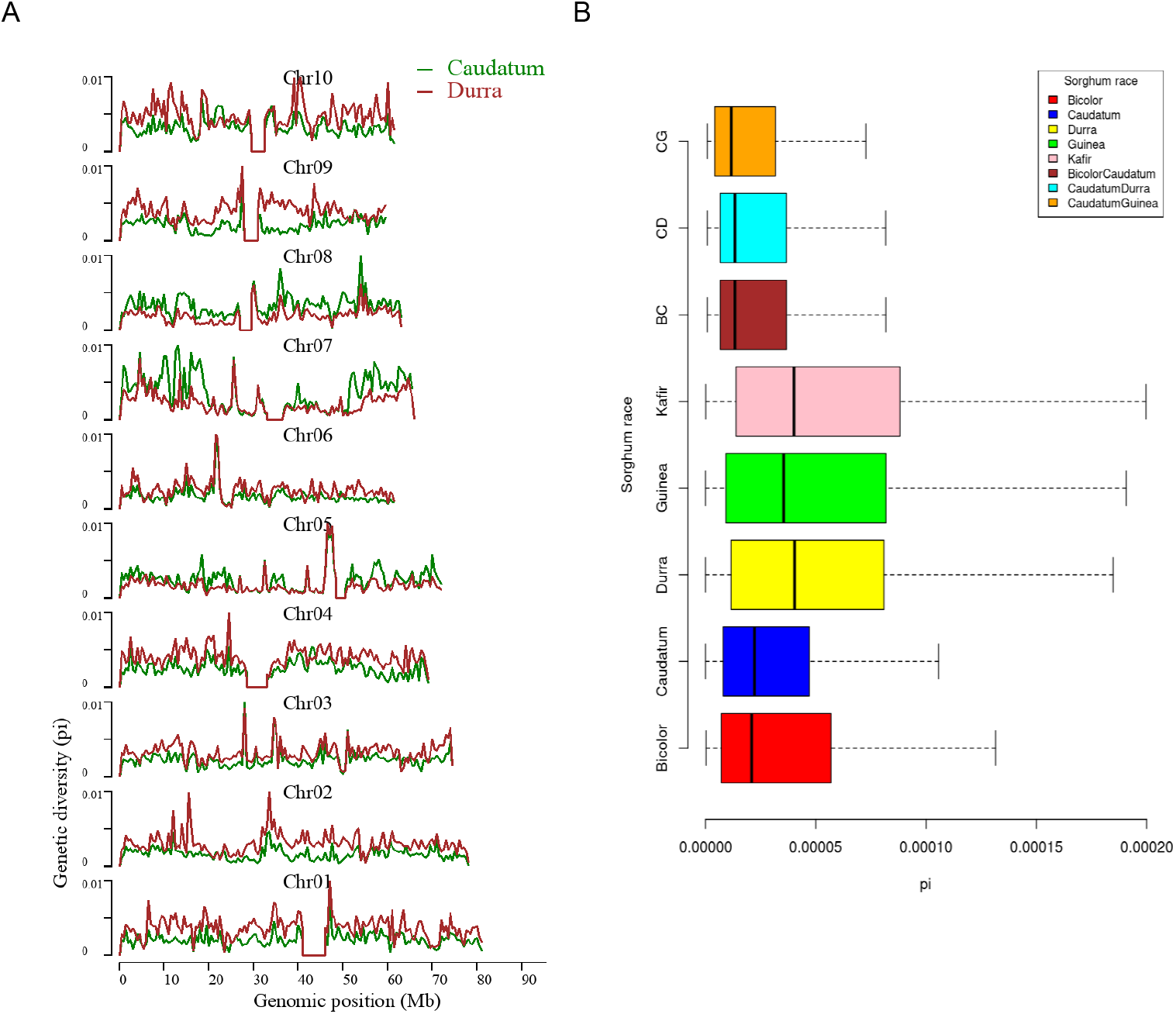
**A.** The genome-wide nucleotide diversity (*π*) density in caudatum and durra populations. **B.** The nucleotide diversity comparison between the five sorghum races and three intermediate races.

We used the *Fst* index to estimate the temporal genetic divergence between the race accessions and observed that the level of genetic differentiation among the sorghum race populations ranged from moderate (*Fst* = 0.044 for caudatum vs durra) to relatively high (*Fst* = 0.18 for bicolor vs guinea) (Supplementary Table 9 and 10; Supplementary Figure 4) indicating that inter-population differences were relatively low. The average *Fst* between the bicolor and other races was ~0.16, which was higher than in non-bicolor race comparisons indicating that gene flow from bicolor to other races was much earlier than the gene flow between the rest (non-bicolor) of the races. The durra and guinea populations revealed the second-highest *Fst* of 0.1228 and were classified as the sorghum race intermediates (Supplementary Table 9). A total of 19,696 SNPs having significant high *Fst* were reported between bicolor-kafir race combinations, of which 910 SNPs were genic SNPs (Supplementary Table 10).

The difference between (diverse) sorghum race populations was measured with Tajima’s D (Table 2). A total of 13,070 SNPs were reported to have *θ_π_* (observed value) less than *θ_k_* (expected value) (maximum 4,612 and minimum 1,869 SNPs from durra and bicolor respectively), indicating that the variants may have undergone a recent selective sweep. Another 311,045 SNPs reported greater *θ_π_* compared to *θ_k_* (maximum 202,684 and minimum 76,836 SNPs from guinea and bicolor, respectively) suggesting a balancing selection. Compared to non-bicolor race mutations, a lower number of mutations were linked to swept genes than with balancing selection genes in the bicolor race (Supplementary Table 11 and 12).

**Table 2.** Summary SNP statistics in Sorghum race populations.

| Race | SNP | $\pi$ ($10^{-5}$) | Tajima's D ($\theta_\pi > \theta_k$) | Non-synonymous SNPs | Synonymous SNPs | Non-synonymous/Synonymous |
| --- | --- | --- | --- | --- | --- | --- |
| Bicolor | 374,545 | 3.37 | 0.44 | 11,100 | 9,957 | 1.114 |
| Caudatum | 961,644 | 2.72 | 0.27 | 31,391 | 31,168 | 1.007 |
| Guinea | 976,558 | 4.30 | 0.37 | 31,827 | 31,681 | 1.004 |
| Durra | 1,261,078 | 3.89 | 0.18 | 38,362 | 37,651 | 1.018 |
| Kafir | 886,880 | 4.72 | 0.26 | 27,779 | 27,458 | 1.011 |

### *K-mer* based divergence

The *k-mer* genetic distance between the sorghum accessions was computed from the size-reduced sketches and distance function developed in the mash tool (Supplementary Table 13). The durra race was the most distinct from the reference pan-genome (Ruperao *et al*., 2021) based on the mean distance of accessions, followed by guinea (Figure 4.A, box plot). The bicolor race was the most closely related race to the reference (Figure 4, box plot). Accession from each sorghum race, SCIV4, PI285039, PI276823, PI665088 and PI665108 from bicolor, caudatum, durra, guinea and kafir, respectively were more genetically distinct from the reference (Supplementary Table 13) and representative of the specific race and therefore used for *k-mer* analysis. These distinct sorghum accessions were in agreement with the NJ distance between the accessions (Supplementary Figure 2).

**Figure 4.**
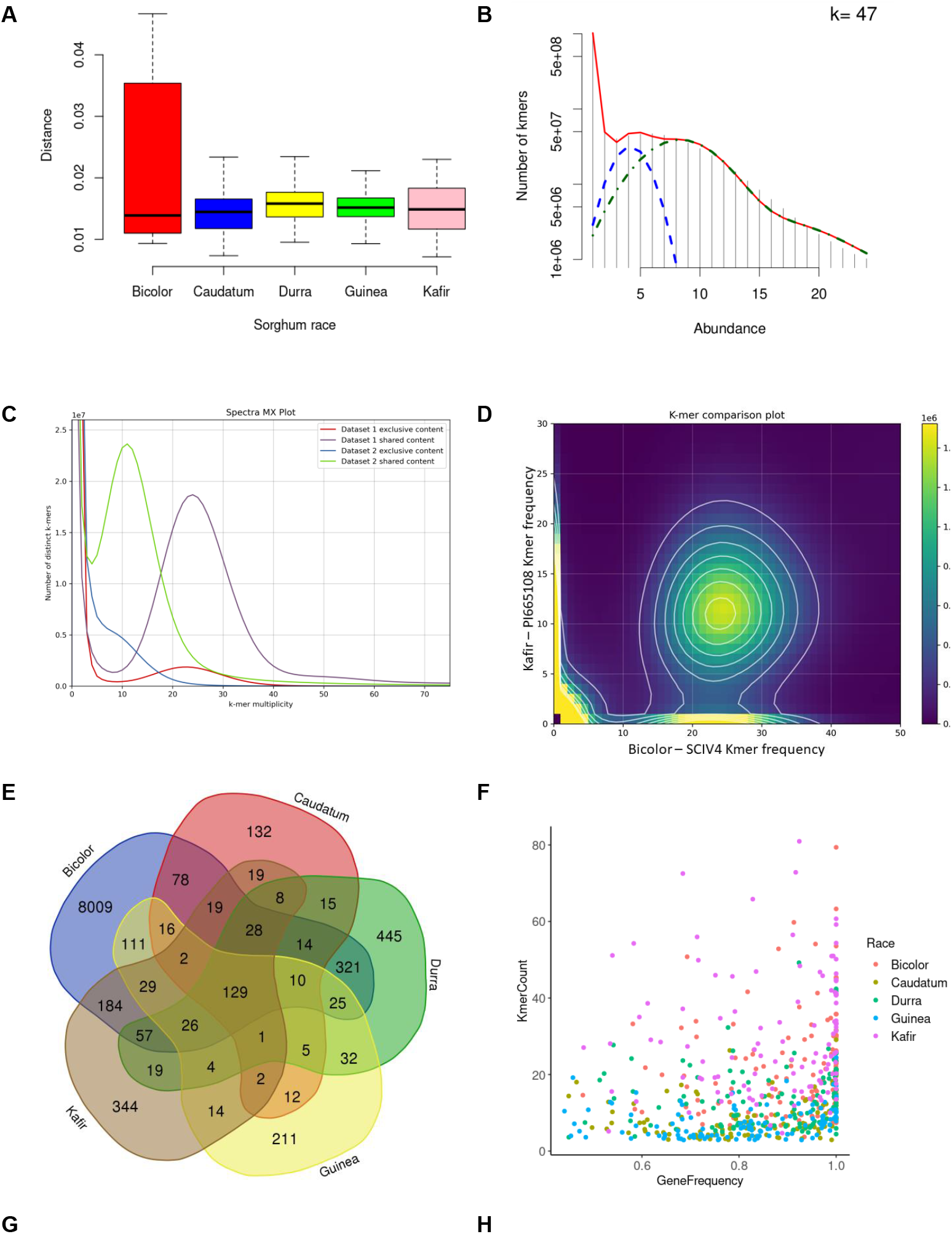
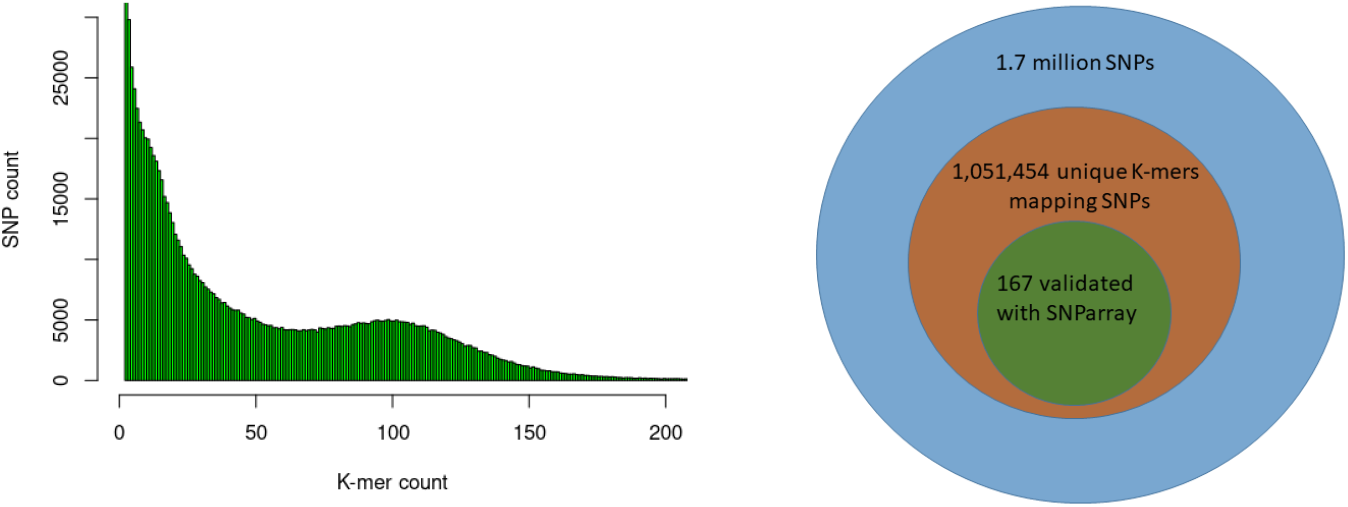
*K-mer* and read mapping overview. **A.** An alignment-free method, the Jaccard index uses the hash procedure to measure the distance between sorghum race accessions. **B** Sampled histogram and fit for 47 *k-mer* lengths. Red is the fit of the complete statistical model of the histogram, blue is the heterozygous *k-mers* and green is the only homozygous *k-mers* **C.** The *k-mers* share between kafir (PI665108) and bicolor (SCIV4) sorghum race accessions as dataset1 and dataset2 respectively. **D.** Distinct *k-mer* share between kafir and sorghum race, the cloud indicates the shared *k-mers* and heigh density *k-mers* on x and y-axis are unique *k-mers* respectively. **E.** Mapped *k-mer* sequence reads in the number of genes in each sorghum race accessions **F.** Proportion of sorghum race common genes covered with horizontal and vertical coverage with *k-mer* sequence reads. **G.** The unique *k-mer*s holding the deepvariant SNPs with respective *k-mer* count and off these, **H.** the proportion of deepvariant SNPs validated.

With the optimized 47 *k-mer* size (Figure 4.B), the overall *k-mer* sequence comparison between the five race accessions (2.3 billion *k-mers*) showed that 35.3% (434 million unique *k-mers*) of common *k-mers* present in all five races accessions, this indicates the conserved *k-mer* of all sorghum race accessions. The 13.3% (314 million *k-mers*) were commonly seen in any four sorghum race accessions, indicating that these *k-mers* were absent in at least any one of the sorghum races. This variability decreased to 8.8% (108 million *k-mers*) and 6.3% (78 million *k-mers*) on measuring the common *k-mers* between three and two sorghum race accessions respectively. For example, SCIV4 (bicolor) and PI665108 (kafir) shared 402 million distinct *k-mers*, which was 45% and 23.5% of total distinct *k-mers* reported respectively (Figure 4.C - D). From this *k-mer* comparison between the sorghum race accessions, 23.8% of *k-mers* were unique to sorghum races. These race-specific *k-mers* were possibly unique to genomic sequence (as a single genome sequence for each race was used for the analysis).

Overall, 10,321 gene PAVs were identified based on the *k-mer* sequence reads mapping to sorghum pan-genome assembly (Supplementary Table 14) (Figure 4.E). The mapping of the race-specific *k-mer* sequence reads identified 490; 9,058; 629; 1,139; and 885 genes in caudatum, bicolor, guinea, durra, and kafir sorghum accessions, respectively. Bicolor had the most unique (9,058) number of genes while caudatum had the least (490). One hundred and twenty-nine (129) genes were commonly present in all sorghum race accessions (Supplementary Table 15), indicating the *k-mer*s are unique with the specific variations or *k-mers* partially mapping the gene length-frequency with horizontal mapping range of 0.4 to 1 (frequency) (Figure 4.F). Furthermore, 1,051,453 SNP were identified supporting the *k-mers* sequence (Figure 4.G) reads of which, 85,048 SNPs were genic and 167 SNPs were validated with the SNParray sequences (Figure 4.H) (Supplementary Table 16) used for sorghum pangenome analysis (Ruperao *et al*., 2021a).

### Selection signatures

Several sweep regions were detected with iHS (Figure 5.A, B and Supplementary figure 5), of which, 64 were significant (FDR < 0.05) (Supplementary Table 17). The majority of sweeps were reported on chromosome 7 (19 regions) followed by chromosome 4 (17 regions) and chromosome 10 (2 regions) (Supplementary Table 17). The highest number of selective sweep regions were observed in durra (54 regions), followed by caudatum (51), guinea (45), kafir (38) and bicolor (30) (Supplementary Table 18 and 19). A total of 14 selective sweep regions were common in all five sorghum races while 21 regions were uniquely absent in any one sorghum race (Supplementary Table 18). For example, 9 selective sweep regions were reported in four sorghum races but uniquely absent in the bicolor race alone (Supplementary Table 18).

**Figure 5.**
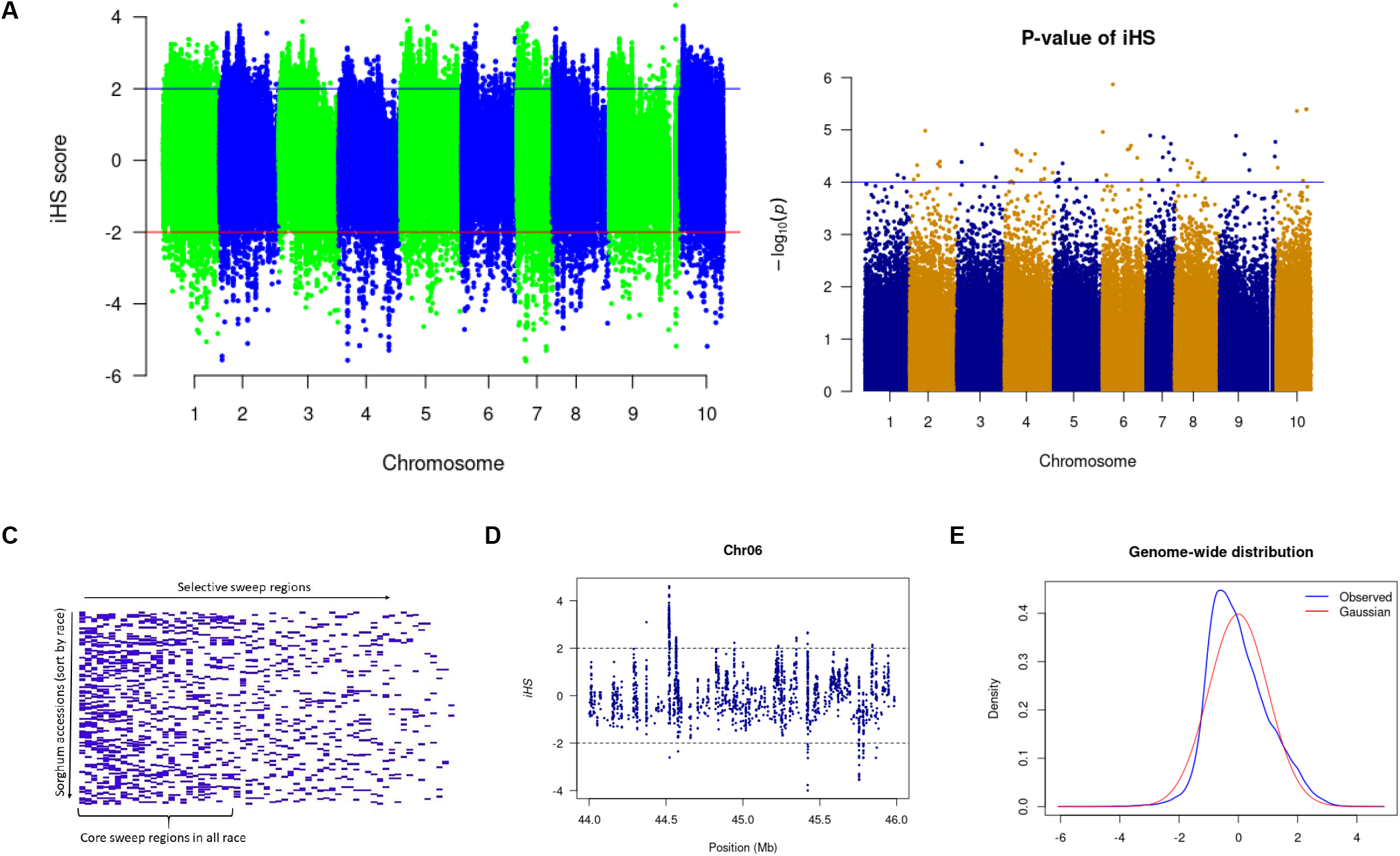
**A)** The iHS score distribution across the genome with **B.** associated p-values. **C.** Selective sweep regions in sorghum accessions sorted by race. **D.** A subset of iHS on chromosome 6 showing the regions having values above and below the threshold value and **E.** The distributions of the standardized iHS scores and comparison with standard Gaussian distribution.

We used the cross-population extended haplotype homozygosity (XP-EHH) score and detected sweep regions from each combination of sorghum race population (Supplementary Figure 5) (Table 3). We identified 8,888 significant (FDR < 0.05) selection sweep regions, of which 3,504 regions were common between more than two sorghum race combinations (Supplementary Table 20-29). Out of all selective sweeps identified from the sorghum race combinations, chromosome 5 had the maximum of 1,399 regions while chromosome 9 had the least (616). The kafir population exhibited the highest (2,473) selective sweep regions in comparison with the guinea race (Supplementary Table 28). Only a few (525) sweep regions were reported in bicolor population (Supplementary Table 23).

**Table 3.** Description of the candidate selective sweep regions detected using XP-EHH between the sorghum race populations

| **Sorghum race combination** | **Significant XP-EHH regions** | **Supporting iHS** | **XP-EHH candidate genes\$** | **XP-EHH mapping genes&** |
| --- | --- | --- | --- | --- |
| Bicolor × Caudatum | 671 | 3 | 212 | 109 |
| Bicolor × Durra | 640 | 3 | 183 | 127 |
| Bicolor × Guinea | 612 | 5 | 201 | 127 |
| Bicolor × Kafir | 522 | 0 | 117 | 61 |
| Durra × Caudatum | 1,832 | 1 | 587 | 414 |
| Durra × Guinea | 2,350 | 4 | 677 | 367 |
| Durra × Kafir | 1,761 | 3 | 424 | 356 |
| Guinea × Caudatum | 2,084 | 6 | 607 | 328 |
| Guinea × Kafir | 2,070 | 1 | 711 | 426 |
| Kafir × Caudatum | 1,263 | 2 | 425 | 365 |

\$ The number of genes refers to the genes mapping 5kb upstream/downstream to selective sweep region. & The selective sweep region present within the gene regions.

### Overlapping selection regions between Tajima’s D and XP-EHH

We defined the overlapping selection regions as those located beyond the thresholds and in the same chromosome sequence location. Tajima’s D statistics were obtained from each sorghum race population dataset and identified the genes which did not fit the neutral theory model at equilibrium between mutation and genetic drift. A total of 324,115 genome-wide bins were observed with non-equilibrium statistics of neutrality test, of which 311,045 (with SNPs in range of 76,836-bicolor to 202,684-guinea) were undergoing purifying selection (negative selection) and 13,070 were (with SNPs in range of 1,869-bicolor to 4,612 durra) selection maintained (balanced positive selection) (Figure 6. A). Among the variants undergoing purifying selection, 43,191 bins had a significant low *Fst* index supporting the signature of a recent population expansion (Figure 6. B), of which 14% were from genic regions (Supplementary Table 30). The purifying selection regions had low diversity (Figure 6. D) with reduced allele frequency in the descendant population compared to the ancestral population.

**Figure 6.**
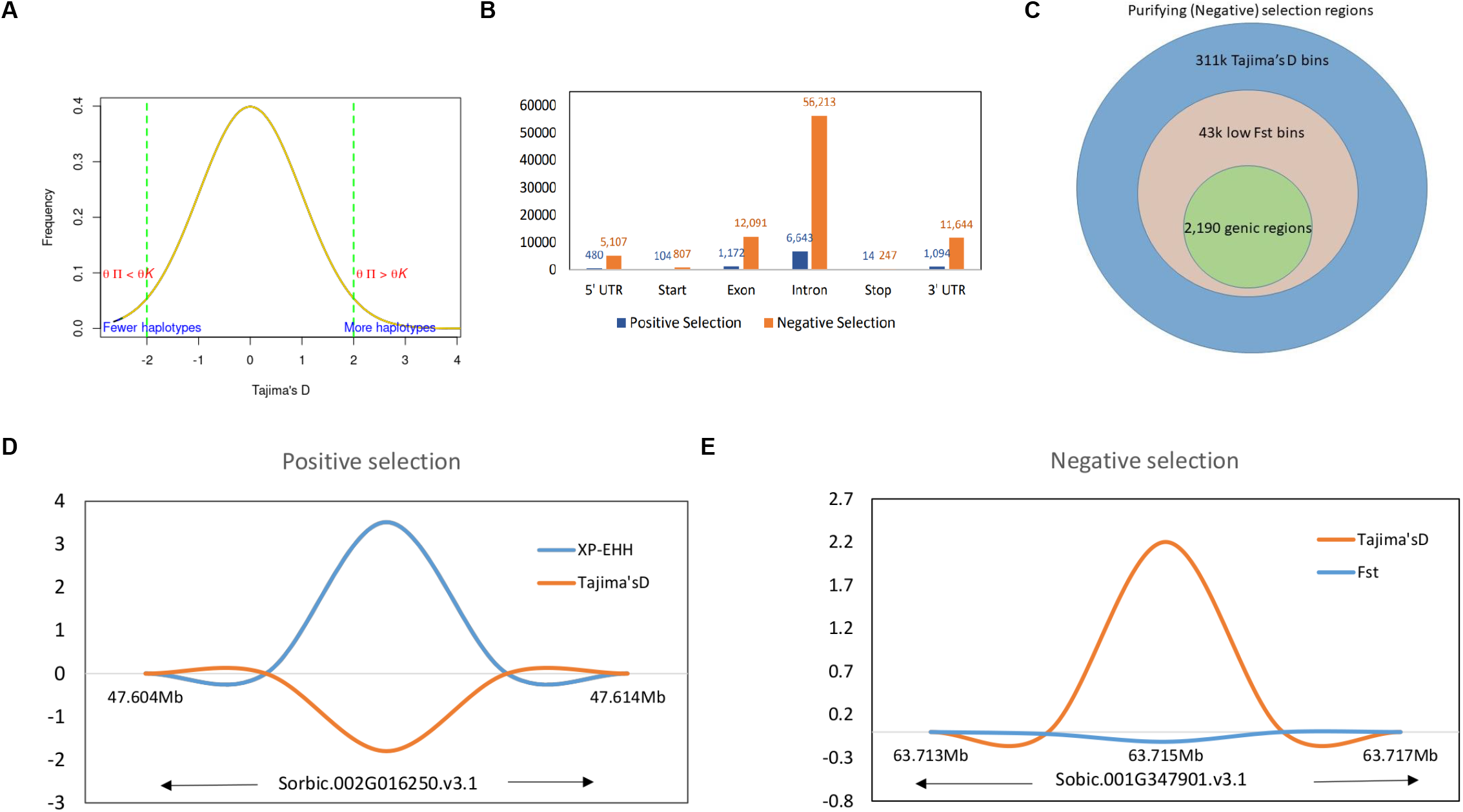
**A.** Tajima’s D values distribution with signifying positive and negative thresholds. **B**. Structural annotations of genomic regions in positive and negative thresholds. **C.** The proportion of higher Tajima’s D values and lower *Fst* valued bins as negative selection regions and corresponding genic regions. **D.** The positive selection region on Chr09 with XP-EHH score and Tajima’s D valued plot and **E.** A genic negative selection region on Chr01 (specific to guinea race) with significantly higher Tajima’s D and lower *Fst* value regions.

The significant selection regions (FDR <0.05) detected by XP-EHH were specific to the pair-wise sorghum race combinations. Among the identified 8,888 significant XP-EHH candidate-sweep regions from overall sorghum race combinations, of which 179 regions were genic (Supplementary Table 31) and “selection-maintained variations” indicating the recent population contraction. The overall sweep regions were in the comparable range identified in other crops such as wheat, (3,105 – 16,141 sweep regions in the genome of domesticated einkorn and emmer lines; Zhou *et al*., 2020) and soybean (3,811 genes positioned within the selective sweep regions)(Saleem *et al*., 2021). Among the chromosomes, chromosome 5 and groups of 4,7,9,10 chromosomes contained the highest (33) and lowest (14) numbers of genes, respectively. A relatively low Tajima’s D was observed in selective sweep regions when compared with a significantly higher XP-EHH valued region (Figure 6. D).

### Enrichment of candidate genes under selection

A total of 2,370 genome-wide genes were observed with non-equilibrium stats of neutrality test, of which 179 were selection-maintained (balanced selection) while 2,191 were undergoing purifying selection (Supplementary table 30 and 31). Durra and bicolor had the maximum (110) and minimum (39) number of genes undergoing positive selection respectively. Bicolor (409) and guinea (1,133) had the maximum and a minimum number of selection-maintained genes.

A similar trend of the least number of genes reported in the bicolor race (421) and guinea has the maximum (1,166) genes under purifying selection. Among the five races, guinea and kafir shares the maximum number of common selection-maintained (26) and purifying (70) genes, suggesting potential rich gene flow between these two races (Figure 7). Additionally, guinea, kafir and durra reported maximum genes as sweeps regions (guinea-kafir: 26, durra-guinea: 23 and durra-kafir: 21) (Figure 7. A), and low nucleotide diversity (*π*) in caudatum and bicolor (Figure 3. B) also indicating the traits regulated by these regions may have undergone similar histories of selection.

**Figure 7.**
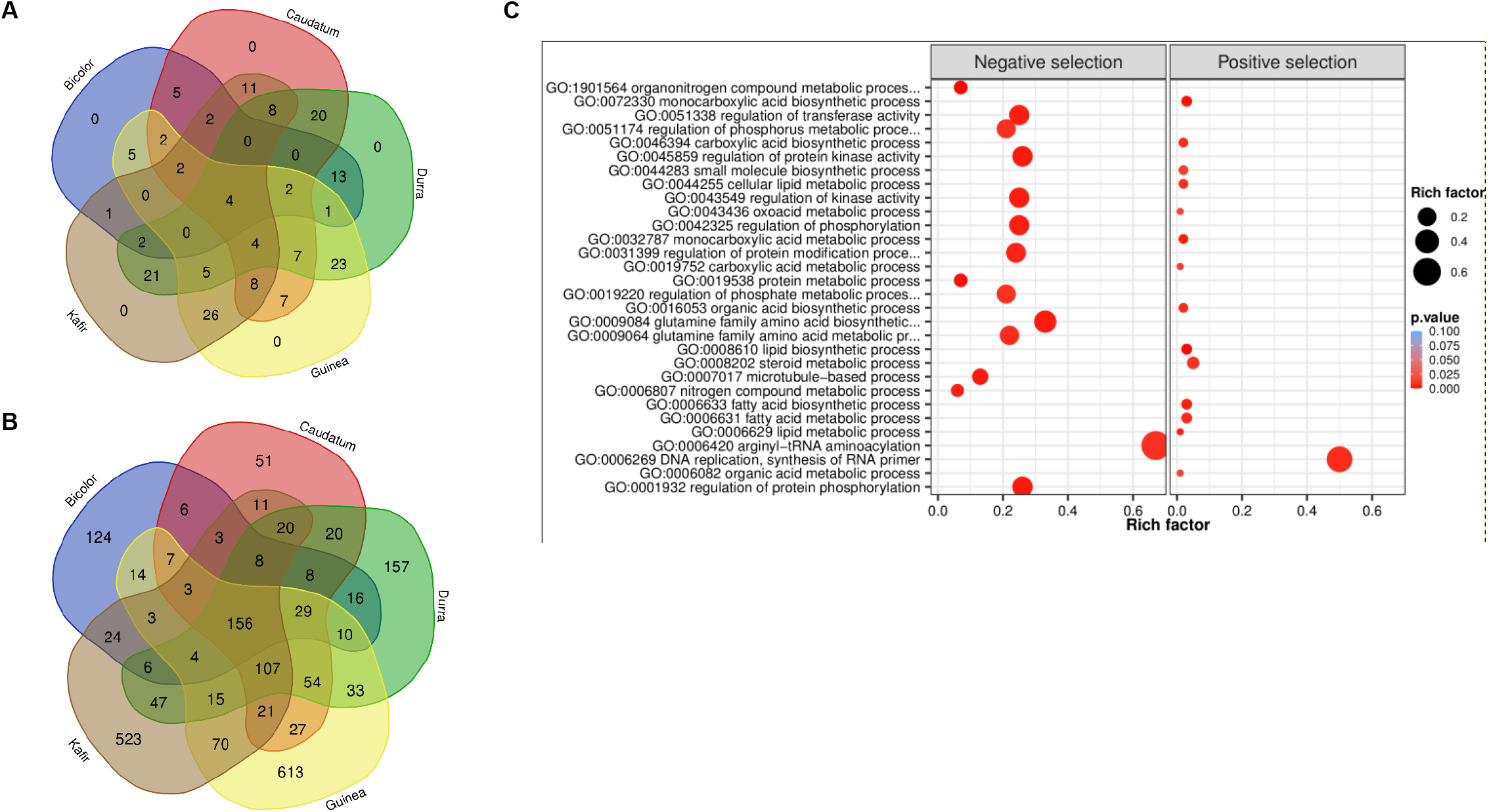
Venn diagram of **A.** positive selection and **B**. negative selection genes from five sorghum races combinations. **C**. Significantly enriched GO biological terms for positive and negative selected genes (bubble color indicates the p-value range and size indicates the gene ratio).

The 2,370 genes undergoing selection pressure (both positive and negative) showed significantly enriched gene ontology (GO) term and among these genes (Figure 7.C, Supplementary Figure 6, 7), the top GO term was lipid biosynthetic process (GO:0008610) and organonitrogen compound metabolic process (GO:1901564) for genes with positive and negatively selection, respectively (Supplementary Table 32). Among the positively selected gene set, most of them were enriched with lipid biosynthetic process (GO:0008610), metabolic process (GO:0006629), carboxylic acid metabolic process (GO:0019752), oxoacid metabolic process (GO:0043436), organic acid metabolic process (GO:0006082), most of these metabolic pathways were related to plant stress resistance. Whereas the negatively selected gene was majorly enriched with nitrogen compound metabolic process (GO:0006807), organonitrogen compound metabolic process (GO:1901564) and protein metabolic process (GO:0019538) (Supplementary Table 32). The nitrogen utilization and metabolic pathway were found significantly enriched and confirmed the genes under selection throughout either domestication or during subsequent breeding with earlier selection study (Massel *et al*., 2016). The genes enriched with ‘DNA replication, ‘lipid metabolism’ and ‘hormone signal’ suggest that sorghum has evolved the defense strategies, and enrichment of phosphorylation, kinase activity, transferase, phosphate and phosphorus metabolic process triggers many metabolic processes and plant growth activity.

Kyoto Encyclopedia of Genes and Genomes (KEGG) pathways were identified according to the selection’s signature candidate gene with a p-value <0.05 (Figure 8). A KEGG pathway enrichment analysis was performed for the selection signature gene to identify the number of significantly changed samples along the pathway that were relevant to the background number. A total of 2,370 genes were mapped onto 315 pathways, and the most enriched sequences were metabolic pathways and biosynthesis pathways. The top 14 pathways with the greatest number of annotated sequences are shown in Supplementary Table 33. Most of the significant pathways were in metabolism, biosynthesis, excision repair and secondary metabolites. The most significantly changed KEGG pathways were resulting in the sphingolipid metabolism (Figure 8.b), betalain, steroid biosynthesis, phosphonate and phosphinate pathways for positive selection genes. Sphingolipids are essential components of plasma membrane providing structural integrity to plant membrane, regulating the cellular process, and also enhancing in improving the tolerance of sorghum to biotic and abiotic stresses. Steroid hormone biosynthesis and the phosphonate and phosphinate metabolism pathways were also involved in the adaptation of sorghum to low salinity. Whereas base and nucleotide excision repair, biosynthesis of secondary metabolites, glycolysis, GPI and nitrogen metabolism were significantly enriched in negative selection genes (Supplementary Table 33). These annotations provide valuable information for studying the specific biological and metabolic processes and functions of genes under selection pressure in sorghum accessions.

**Figure 8.**
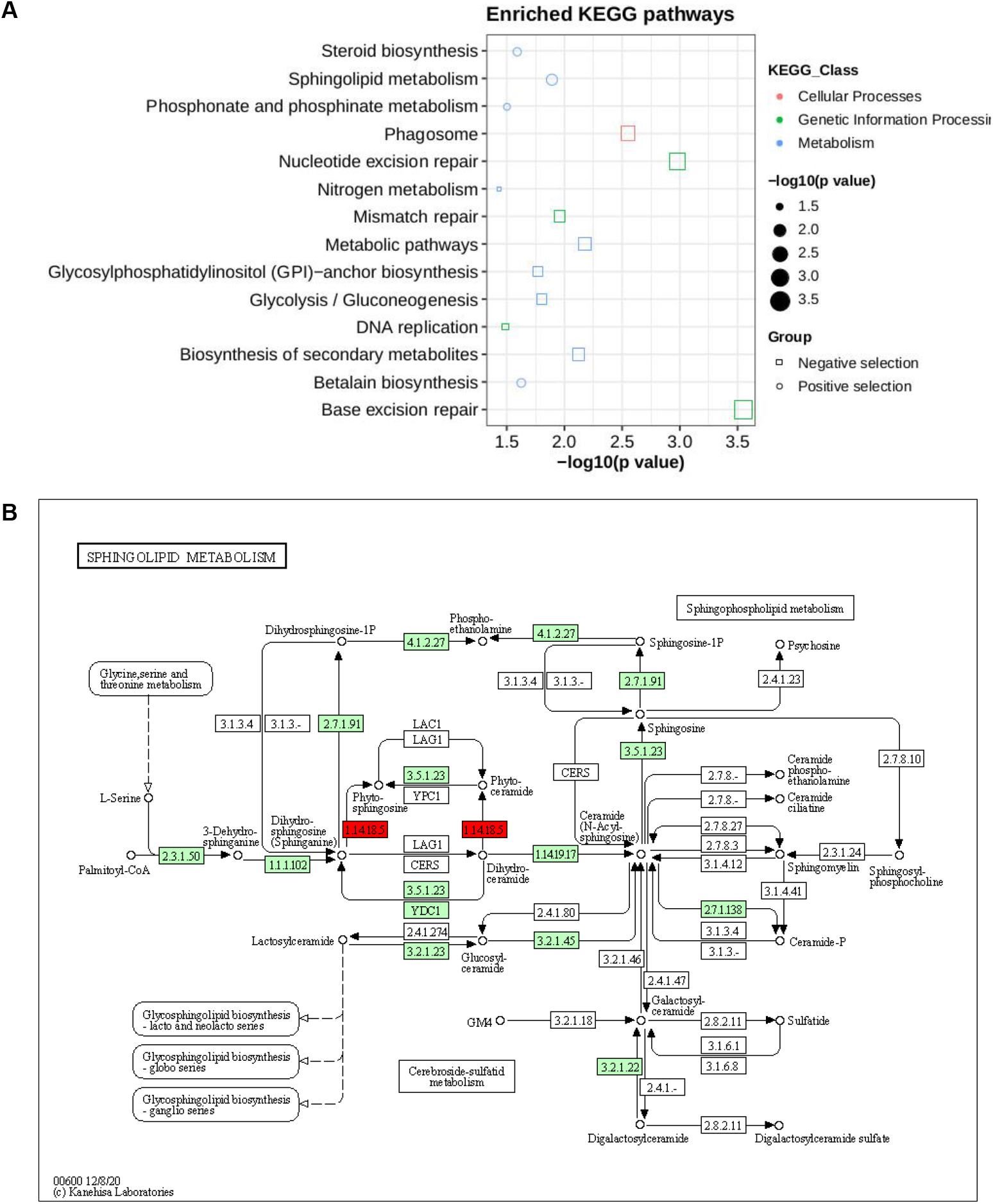
**A.** KEGG pathway enrichment for genes under selection pressure **B.** Pathway of sphingolipid metabolism. The two red boxes represent the positively selected genes involved in pathways.

### Overlap of signatures of selection with QTLs

Quantitative trait loci associated with seven traits that overlapped with detected signatures of selection were compared with earlier reported sorghum QTLs (Hostetler *et al*., 2021). Analysis of the overlaps between signatures of selection and reported QTL indicated that 10 and 206 linked genes were identified as positive and negatively selected genes respectively (Supplementary Table 34, 35). Some QTL for traits of plant height, root biomass, dead above-ground biomass, live above-ground biomass and total biomass overlapped significantly with putative gene regions of signatures of selection.

## Discussion

We have demonstrated the utility of vast sorghum genomic data that exists in public databases for characterisation of a representative set of sorghum (Valluru *et al*., 2019). Our results validate the application of deep learning for the characterisation of sorghum races and goes further to establish nucleotide diversity and genetic divergence across and within different sorghum races. We also used existing QTL data to identify candidate genes that are under both negative and positive selection.

### Germplasm selection

The diversity of sorghum types or races has been related to selection, geographic isolation, and gene flow from wild to cultivated plants (Wet and Huckabay, 1967; HARLAN and STEMLER, 2012). The bicolor race has been considered the progenitor of more derived races based on its primitive morphology (Harlan and Wet, 1972; Stemler *et al*., 1975), while the guinea represents independent domestication from West Africa (Deu *et al*., 2006; Morris *et al*., 2013). The durra race was associated with dryland agricultural areas along the Arabic trade routes from West Africa to India (Wet and Huckabay, 1967). The kafir race was associated with Bantu agricultural tradition while the caudatum is believed to have originated from central and eastern Africa.

The sorghum reference set used in the current study was earlier selected by Billot *et al*. (2013) after genotyping 3,367 global collections using 41 representative nuclear SSR markers and is considered to be representative of the sorghum collections that exist in various global gene banks. Our results confirm that the clustering of sorghum germplasm was largely according to regions (Paterson *et al*., 2009; Bekele *et al*., 2013; Ramu *et al*., 2013) indicating continuous gene flow between various racial groups depending on where the sorghum races are grown. The only distinct exception was in the guinea race, which was expected since the guinea race is specifically grown in West Africa and therefore any gene flow would be confined within the West African locations.

Our study also identified many intermediate accessions (more than 15 accessions) as a result of the continuous gene flow suggesting that a different criterion other than morphology will need to be used in future studies for the correct classification of sorghum races and their intermediates.

Capturing race-specific sequences will be critical in future studies for the follow-up identifications of variants and/or, genes associated with each sorghum race. For example, longer k-mers (>15 bp) have been utilized as biomarkers (Drouin *et al*., 2016; Wang *et al*., 2018) as they can hold biological information and depict specific signatures in nucleotide sequences (Wang *et al*., 2016). Our ability to differentiate abundant k-mers between the different sorghum races in the current study provides an opportunity for future studies to utilize k-mers as race- or accession-specific identifiers in sorghum.

### DeepVariant calling and its utility in sorghum breeding

For the first time in sorghum, we used the DeepVariant (Poplin *et al*., 2018c) tool, a deep learning approach for SNP calling, and reported over two million genome-wide variants from existing sequencing data. One of the concerns of SNP calling from NGS data is the accuracy of SNPs. A recent comparison of SNPs called from the traditional SNP calling tools such as GATK (DePristo *et al*., 2011) with DeepVariant method reported superior performance of the latter (Lin *et al*., 2022), further validating our choice to implement this method in sorghum. Our results were largely consistent with previous studies in sorghum that involved SNP calling from NGS data, including the patterns of SNP distribution observed across the genome (Paterson *et al*., 2009; Bekele *et al*., 2013) and the non-synonymous to synonymous SNP substitution ratio. Our results were also within the range reported for other genome-wide studies such as in soybean (Lam *et al*., 2010), rice 1.2 (McNally *et al*., 2009), and Arabidopsis 0.8 (Clark *et al*., 2007).

Our study made use of short Illumina reads, which are still the most abundant in public databases for sorghum. However, there are now deep learning pipelines available for long reads (Shafin *et al*., 2022) that will need to be considered appropriately when long reads become available in sorghum. Future studies will need to compare DeepVariant with other existing methods and validate our results in different germplasm sets, such as the sorghum diversity panel (Casa *et al*., 2008). Such future studies will also need to pay special attention to sequence coverage and how it would affect the accuracy of variants called. Our study used a minimum overall coverage of 5x, which was more than adequate even for a less efficient SNP calling pipeline (Wu *et al*., 2019). Sequence coverage is one of the major factors affecting the accuracy of SNPs called from NGS datasets, especially in heterozygous species (Gong and Han, 2022). A coverage of 0.01x has been reported as the most cost-effective coverage in sorghum, with 94.1% SNP accuracy (Jensen *et al*., 2020). There will be a need for additional studies establishing the effect of various levels of coverage in the NGS datasets for DeepVariant calling, and how it would affect the SNP accuracy in sorghum.

### Nucleotide diversity and divergence in sorghum

Our study was purely based on existing data and did not allow for much flexibility in the number of genotypes per race. The overall nucleotide diversity observed for sorghum of *π* = 0.000048 is significantly smaller than previously reported by Faye *et al*. (2019) but comparable to a more recent study (Sapkota et al, 2020) that reported *π* = 0.000032. This figure is much lower than for other cereals such as wheat (*π*_A_=0.0017, *π*_B_ = 0.0025 and *π*_D_ = 0.0002) (Zhou *et al*., 2020), maize (*π* = 0.014) (Tenaillon *et al*., 2001) and rice (*π* = 0.0024; Huang *et al*., 2010) and could be a consequence of the few genotypes used in the study. The race-specific nucleotide diversity indicated that the caudatum (57 genotypes; *π*_C_ = 0.0000419) had the lowest diversity followed by guinea (68 genotypes; 0.0000631) and durra (82 genotypes; *π*_D_ = 0.0000637) races. This is in contrast with earlier studies that reported, the guinea race as the most genetically diverse sorghum type (Morris *et al*., 2013; Faye *et al*., 2019). Comparing our results and those of Morris *et al*. (2013) and Faye *et al*. (2019), suggests a positive correlation between the number of genotypes per race with the nucleotide diversity. More studies need to be done to confirm the effective population size per sorghum race that will be optimum for a reliable and consistent nucleotide diversity result.

### Selection signatures

We used two approaches to detect selection sweeps across the sorghum genome, both of which are haplotype-based. The iHS method, which is based on a single population, was meant to detect recent positive selection (Voight *et al*., 2006), while the XP-EHH is based on the comparison of two populations and is considered powerful in detecting beneficial alleles shortly before, or at fixation (Alexandra *et al*., 2015). Such multiple statistical approaches were earlier used for selection sweeps in other crops like cotton (*Gossypium herbaceum*) (Nazir *et al*., 2020) and soybean (*Glycine max*) (Zhong *et al*., 2022). A recent study comparing different methods used for detecting selection sweeps reported that both iHS and XP-EHH were able to identify genomic regions undergoing selective sweep under a wide range of population structure scenarios (Vatsiou *et al*., 2016). Previous studies in sorghum have also reported evidence of selective sweep in sorghum (Casa *et al*., 206; Faye *et al*., 2019) although the methods used for detection were different. Our results on selective sweep regions were further strengthened by Tajima’s D results, which enabled us to identify candidate genes in the significant selective sweep regions.

The 2,370 candidate genes identified in our study (for selection pressure), of which, 7.5% are positively selected, are similar to the proportion of genes identified for domestication and improvement using the gene-based population study by Mace *et al*., (2013). The genomic regions that are either positively or negatively selected in the respective sorghum races would give a hint on geographic preferences. More studies will need to delve deeper into specific regional selection sweeps that could eventually be used to predict ideal phenotypes. The remaining candidate genes that were reported as undergoing negative selection with evidence from both Tajima’s D and Fst index values will need. Such genomic analysis of crop landraces would enhance our understanding of the basis of local adaptions (Li *et al*., 2017; Swarts *et al*., 2017).

Some of the trait-associated genes undergoing the selection pressure that have been reported include the dry pithy stem gene mutation that led to the origin of sweet sorghum (Zhang *et al*., 2018), local adaptation to parasite pressure and signatures of balancing selection surrounding low germination stimulant (Bellis *et al*., 2020) and, the strong selection pressure on the sorghum maturity gene (Ma3) (Wang *et al*., 2015). Comparative population genomics assist to dissect the domestication and genome-wide effects of selection as studied in cotton reports 311 selection sweep regions associated with domestication and improvement (Nazir *et al*., 2020) and the selection sweeps identified in the wild and domesticated soybean accessions (Zhong *et al*., 2022).

The population subjected to strong selection pressure may experience the bottleneck and result in a loss of genetic diversity. The level of diversity preserved in a population depends on the background of the emerging adaptive alleles (Wilson *et al*., 2017). Identification of such a large number of selection sweeps suggests the domestication bottlenecks. The identified selection sweeps overlapped with highly differentiated regions suggesting the occurrence of differentiation due to human-mediated selection. These regions help in understanding the genetic basis of domestication and improvement in traits. On further comparison of the selection regions with significant loci of GWAS analysis (narrow down the region), to determine the genes concerning domestication and selection in the sorghum crop.

The results from this study can lead to understanding the changes at the genomic level caused by domestication, selection and improvement of sorghum accessions.

## Methods

### Plant material

We used 272 sorghum accessions, which included accessions that had been used in a previous sorghum pangenome study (Ruperao *et al*., 2021b) and six sorghum bicolor accessions reported in Yan *et al*. (2018) (Supplementary Table 1). Among these genotypes, 82, 21, 57, 68, 21 were durra, kafir, caudatum, guinea and bicolor respectively, while the remaining were mixed accessions.

### Variant discovery

The fastq sequence reads generated from the 272 sorghum accessions were trimmed with Trimmomatic 0.39 (Bolger *et al*., 2014). Alignments to the sorghum pangenome (dataverse.icrisat.org, https://doi.org/10.21421/D2/RIO2QM) as a reference (Ruperao *et al*., 2021c) were performed using Bowtie2 version 2.4.2 (Langmead and Salzberg, 2012). All alignments were converted to binary files with Samtools 1.13 (Li *et al*., 2009) followed by filtering out the read duplication with Picard tools (http://broadinstitute.github.io/picard). The open-source DeepVariant (https://github.com/google/deepvariant) (Poplin *et al*., 2018c) tool was used to create individual genome call sets, followed by merging call sets with Bcftools 1.9 (Bcftools by samtools) then analyzing the merged call set. The merged variants were filtered with ‘maf 0.01 min-meanDP 2 minQ 20’. Filtering was done using Vcftools 0.1.16 (Danecek *et al*., 2011). Retained high-quality sites were used for downstream analysis. Functional annotation of SNPs was done using SnpEff v.4.3 (Cingolani *et al*., 2012).

### Counting *k-mers*

The *k-mer-based* genetic distance between 272 sorghum accessions was measured with Mash (Ondov *et al*., 2016). Out of the 272 accessions used, the mean distance values within each race were used to compare *k-mer*s between the sorghum races. To compare sequences across sorghum races, we determined *k-mer* frequency in sequencing reads from all samples. To identify the common and unique genomic sequences between the sorghum races, we split the sequencing reads into *k* length of the sequence. The optimal *k-mer* size for identifying the distinct *k-mers* was estimated using KmerGenie (Chikhi and Medvedev, 2014) within the *k* range of 21 to 121. The optimized *k*=47 was used for measuring the *k-mer* frequency as shown in the Figure 9.

**Figure 9:**
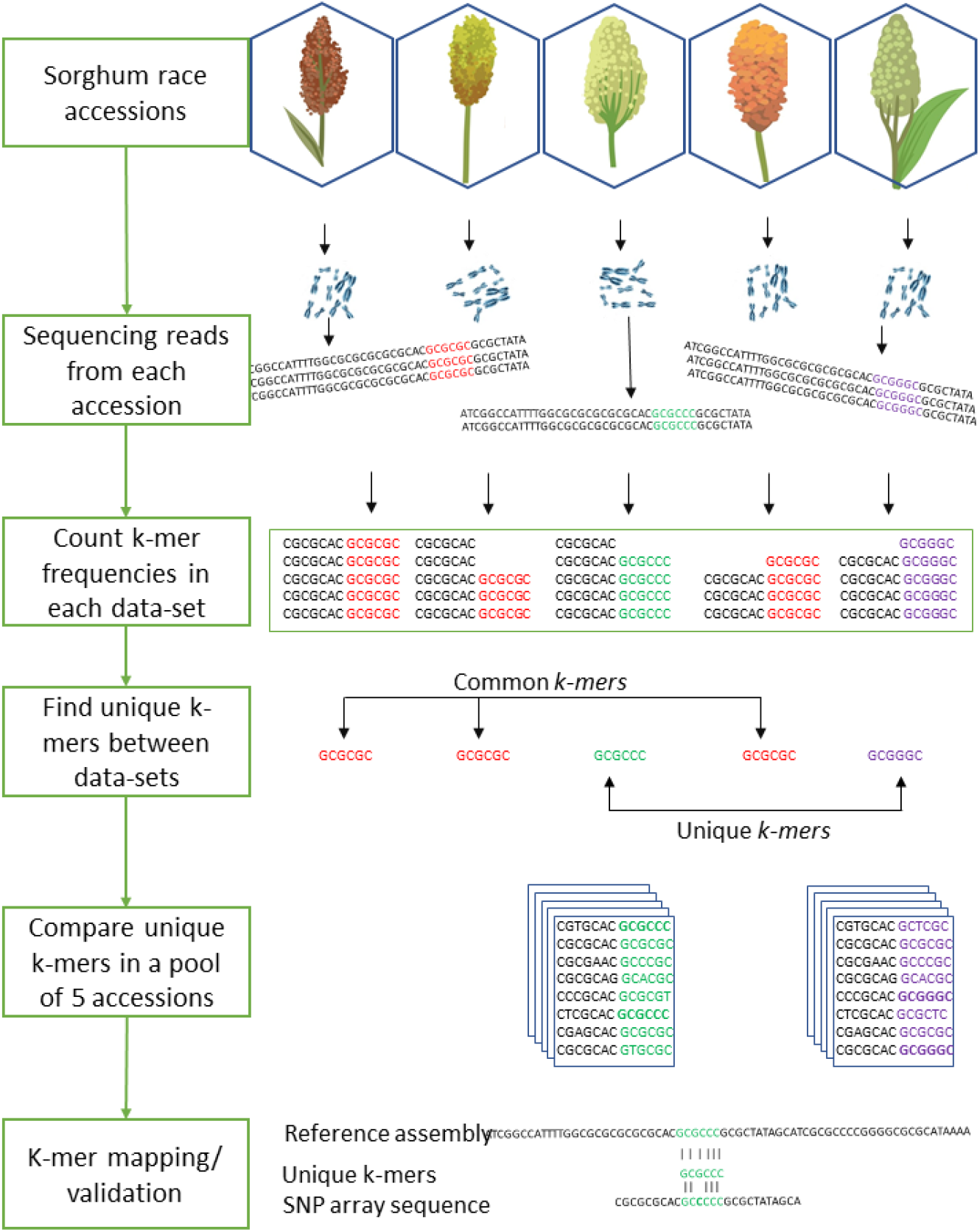
The workflows of the sorghum race accessions comparison with *k-mer* analysis.

We used the hash-based tool Jellyfish (Marçais and Kingsford, 2011) to count *k-mer*s with the optimized *k-mer* length of 31 and filtered out *k-mers* that appeared only once in samples as they were likely from sequencing errors. The *k-mer* hashes were visually inspected through KAT density plots (Mapleson *et al*., 2017) for all five sorghum race accessions by producing the *k-mer* frequency, GC plots and contamination checks. Unique *k-mer*s mapped to sorghum pangenome were validated with mapped SNParray region from a previous study (Ruperao *et al*., 2021b). Based on the mapped *k-mers*, the gene PAVs were extracted from the sorghum pan-genome genes PAVs catalog (Ruperao *et al*., 2021b) with in-house developed script.

### Nucleotide diversity and relatedness

The filtered SNPs were further subgrouped based on race-specific variant alleles. Nucleotide diversity (*pi*) was calculated using Vcftools 0.1.16 (Danecek *et al*., 2011). The *pi* (*π*) distributions were compared to assess changes in genetic diversity over time. The *pi* (*π*) density plots were generated with in-house developed scripts.

In addition, a 1,000 bootstrap resampling was used to estimate the genetic relationship among the accessions with R “ape” (Paradis *et al*., 2004) package to construct a NJ tree and visualized it in iTOL tree viewer (Letunic and Bork, 2019).

### Population differentiation and signatures of selection

Tajima’s D (Tajima, 1989) and per-site *Fst* (based on Weir and Cockerham’s *Fst* estimator) (Weir and Cockerham, 1984) were calculated using Vcftools 0.1.16 software (Danecek *et al*., 2011). Integrated haplotype score (iHS) (Voight *et al*., 2006) analysis was performed using the “rehh” package (Gautier *et al*., 2017) in R v 3.6.3, while the extended haplotype-based homozygosity score test) (XP-EHH) (Sabeti *et al*., 2007) was derived using Beagle (Browning *et al*., 2021). Significant selective sweeps were detected using the Bonferroni FDR threshold (*P* < 0.05).

Overlap of putative genomic regions under selection with previously known QTLs was detected after downloading the mapped QTL regions from Hostetler *et al*. (2021) and comparing them with the identified selection regions.

## Conclusions

This study compared the genomes of the sorghum races with short *k-mer* length sequence to identify the conserved and signature patterns of sorghum race sequences. We implemented a deep learning method to detect the variants and compared structural and functional annotations. On applying the *k-mer*-based genome comparison among the sorghum races, we were able to identify the unique *k-mer* sequences that is specific to the sorghum races and also possibly use as race-specific or accession specific (if *k-mers* compared between accessions) genetic markers. Our study observed a relatively lower genetic diversity in the *caudatum* and *bicolor* races than in *kafir, guinea* and *durra* races. Our results revealed several putative footprints of selection that harbor interesting candidate genes associated with agronomically important traits using different statistical approaches. The findings will enhance our understanding of the dynamics of the sorghum race genomes and help to design strategies to breed better genotypes.

## Author contributions

PR conceived the research. PR, PG, MS and RRD carried out the research. SS managed the computational resource. SPD and DAO provided inputs on sorghum history and domestication. NT, EH, EM and BN assessed and provided inputs for manuscript. PR, NT, AR and DAO drafted and edited the manuscript. All authors read and contributed to the manuscript.

## Acknowledgments

The authors also acknowledge the supporting funds from AVISA (OPP1198373) and ICAR-BMGF (101165).

