## Supplementary Figures1-6 for "DeepVariant calling provides insights into race diversity and its implication for sorghum breeding"

A

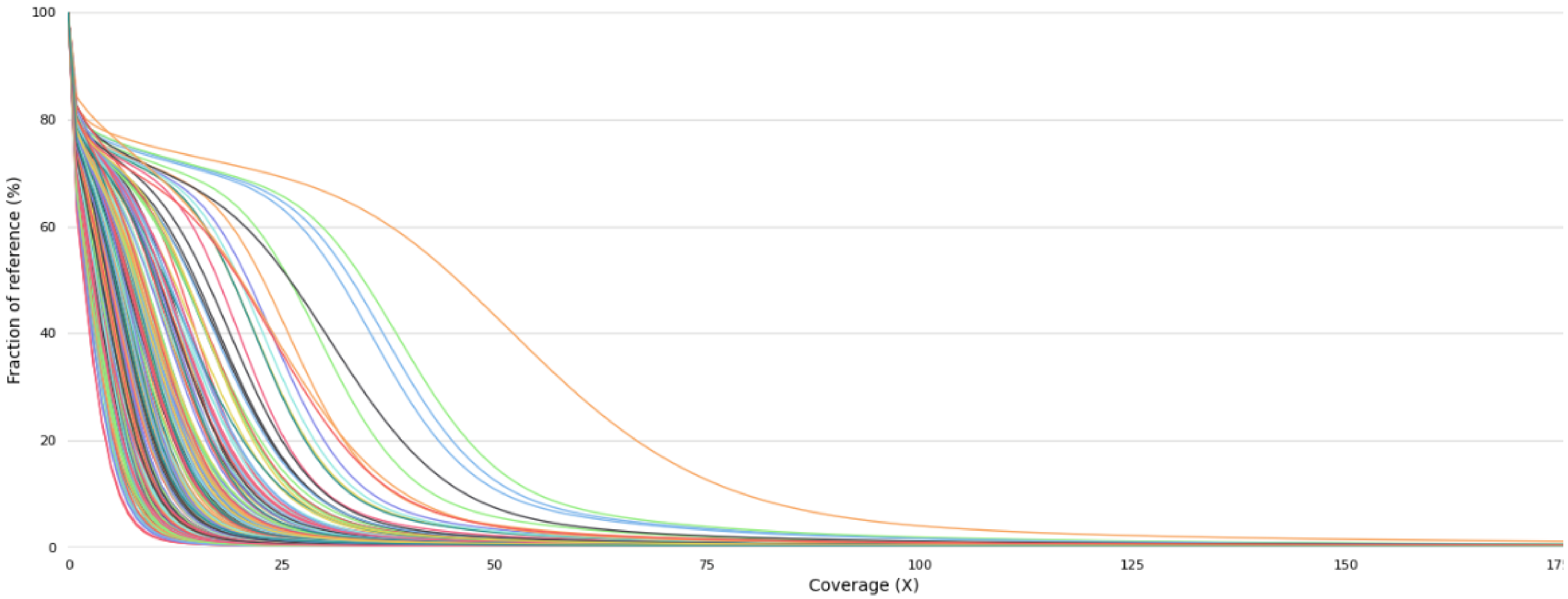

100
80
60
40
20
0
Fraction of reference (%)
0
25
50
75
100
125
150
Coverage (X)

B

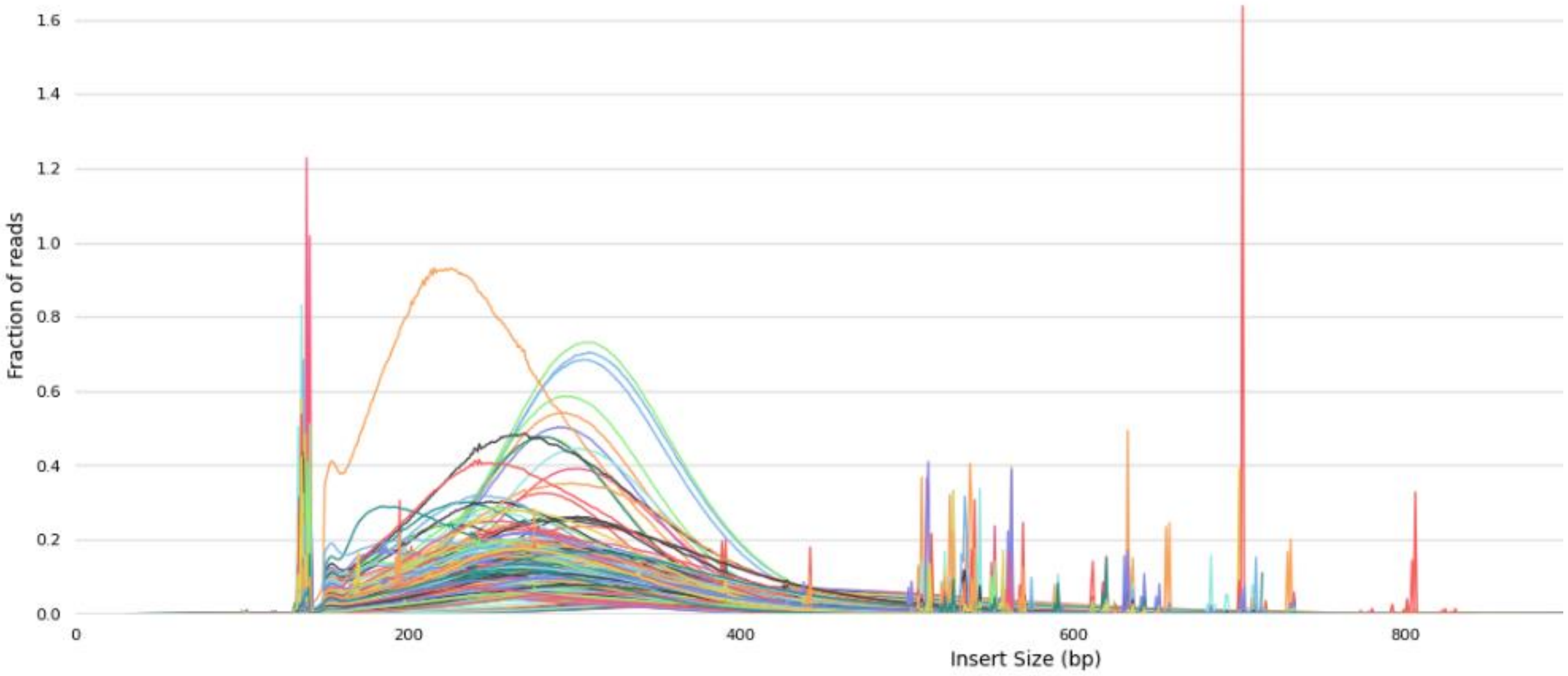

1.6
1.4
1.2
1.0
0.8
0.6
0.4
0.2
0.0
Fraction of reads
0
200
400
600
800
Insert Size (bp)

C

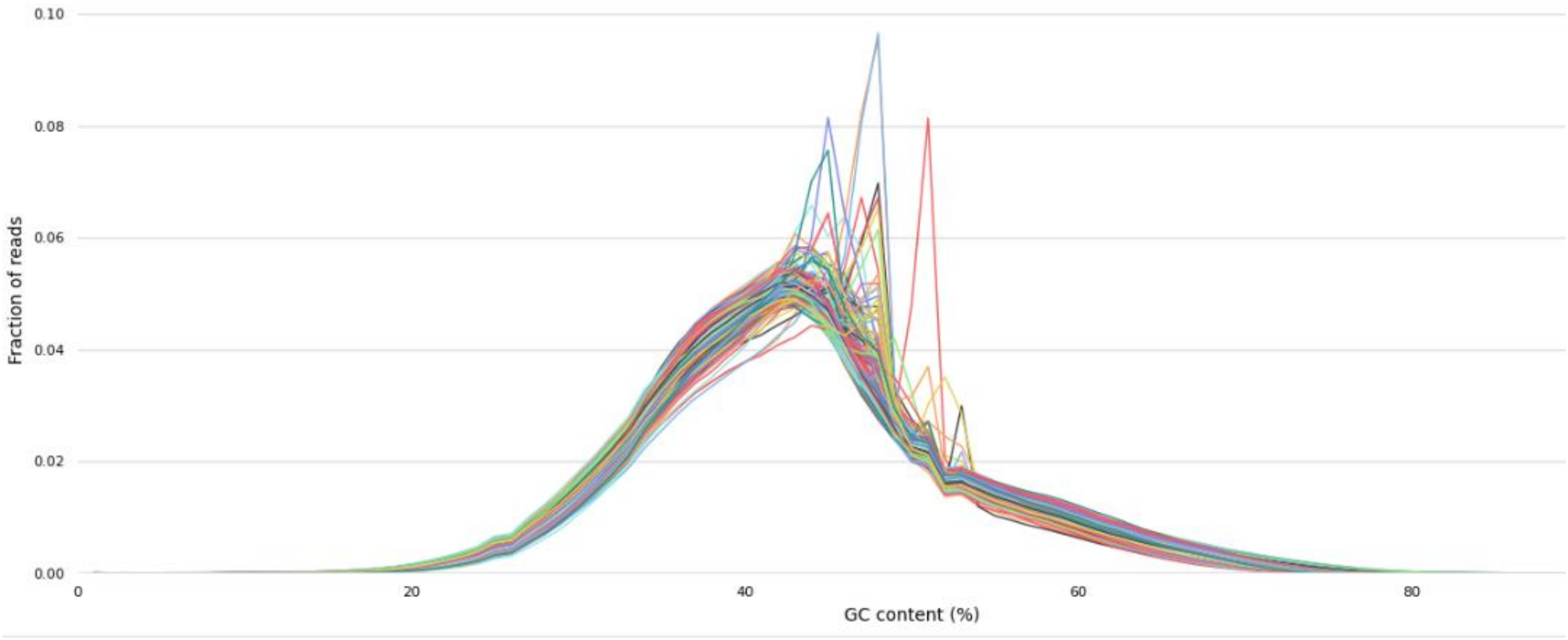


**Supplementary Figure 1:** Sorghum WGRS mapped stats showing the **A)** reference genome covered in both horizontal and vertical coverage, **B)** inset size of complete datasets **C)** GC content in all sorghum accessions.

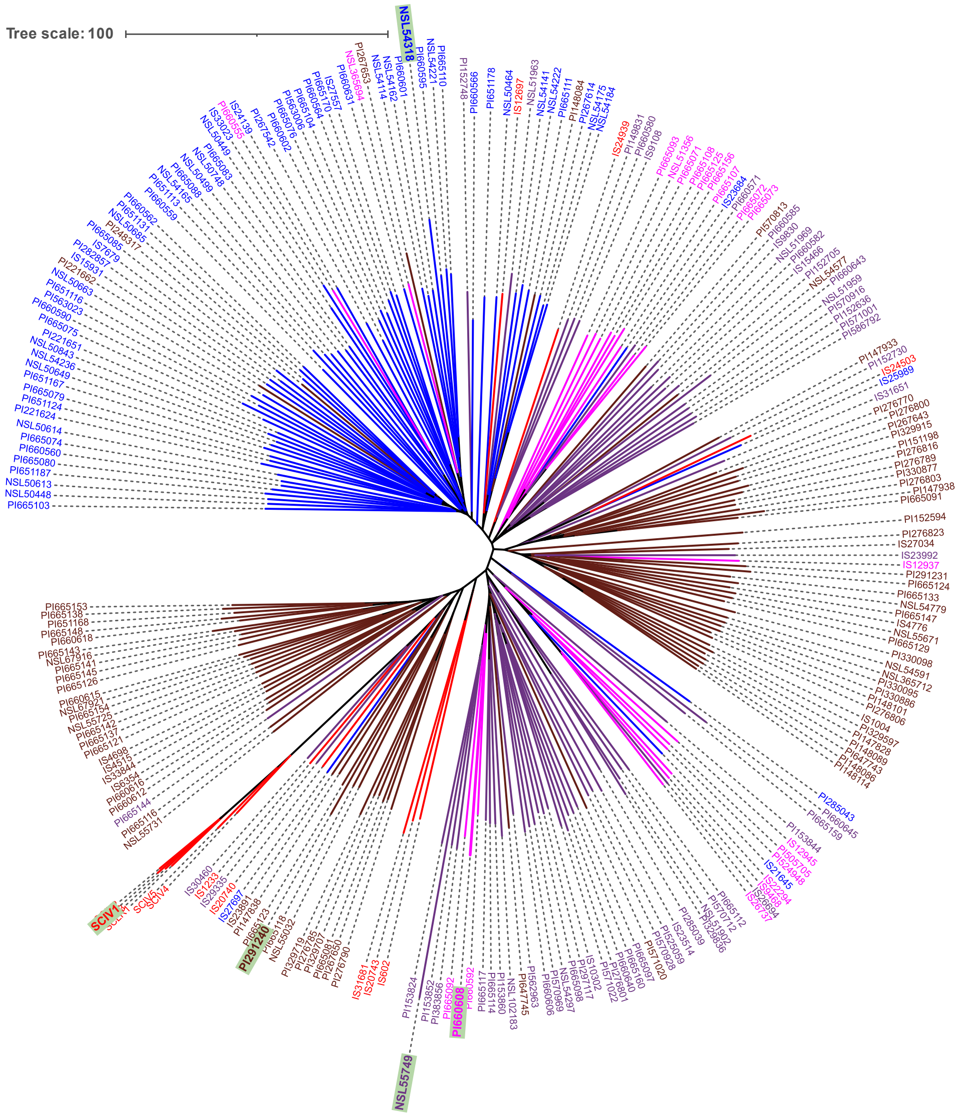


**Supplementary Figure 2.** Sorghum race accession NJ tree with nodes in blue color are *Guinea*, pink colour accessions are *Kafir*, red color for *bicolor*, brown color in *Durra*, purple color for *Caudatum*. Highlighted node labels are the most distinct accessions in respective sorghum race accessions. The phylogenetic tree contained a distinct cluster of 63 *Guinea* race accessions (nodes in blue colour) mixed with a few other race individuals, like PI221662, PI248317, PI267653 and PI148084 from *Durra* race (nodes in brown colour), PI660555 and NSL365694 from *Kafir* race (nodes in pink colour), IS12697 from *bicolor* race (nodes in red colour). The other sorghum race clusters were split with non-corresponding sorghum race accessions. For example, *Durra* race has 91 accessions split into two clusters with *Caudatum* and *Kafir* race accessions. The sorghum *bicolor* race which was assumed to be more distinct to other race were placed mostly in *Durra* and *Guinea* race cluster. Among the *bicolor* race, the China origin accessions were grouped distinctly in *Durra* cluster compared to other *bicolor* race accessions.

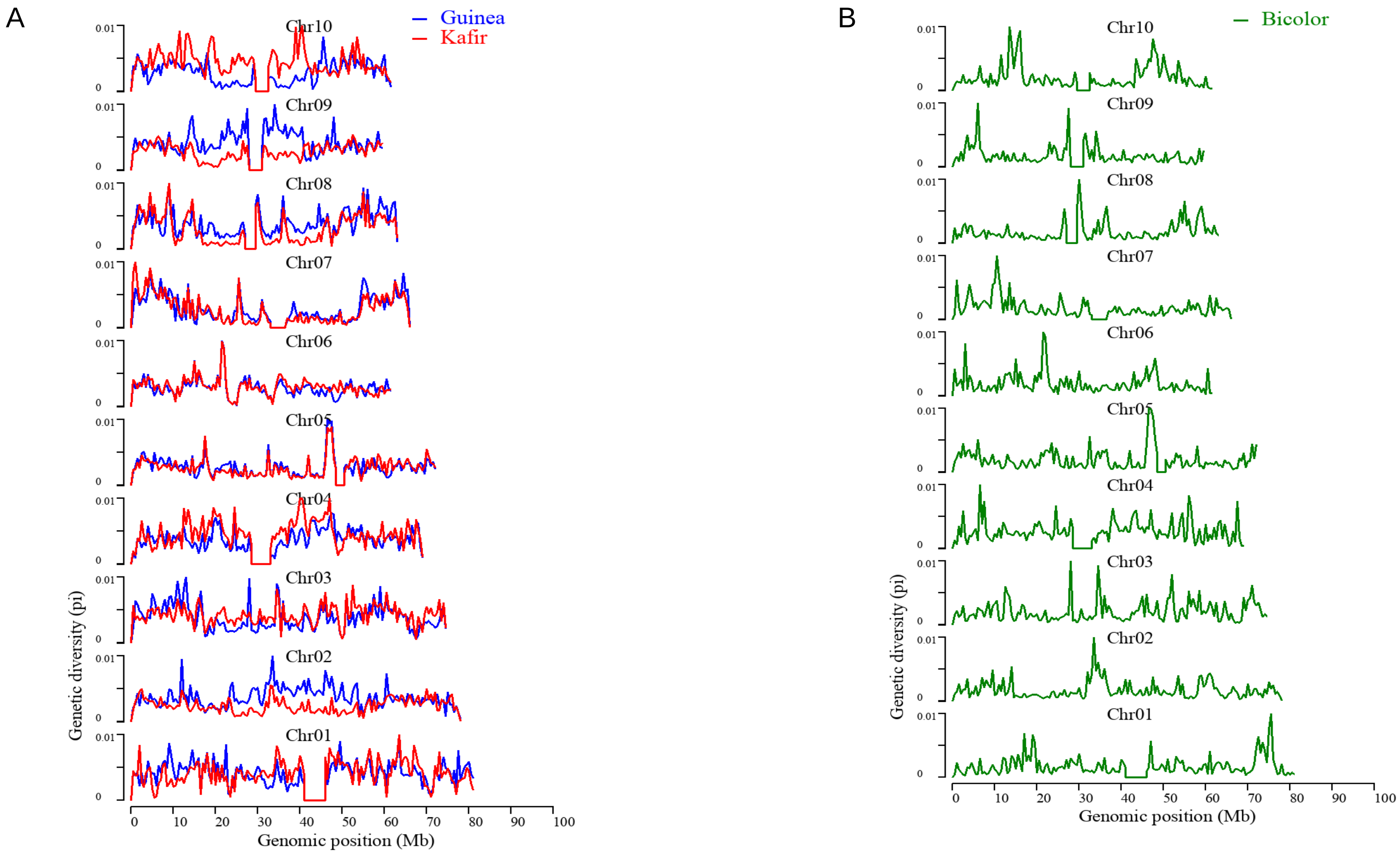


**Supplementary Figure 3.** Nucleotide diversity (π) in sorghum race populations **A)** Guinea and kafir **B)** Bicolor

A

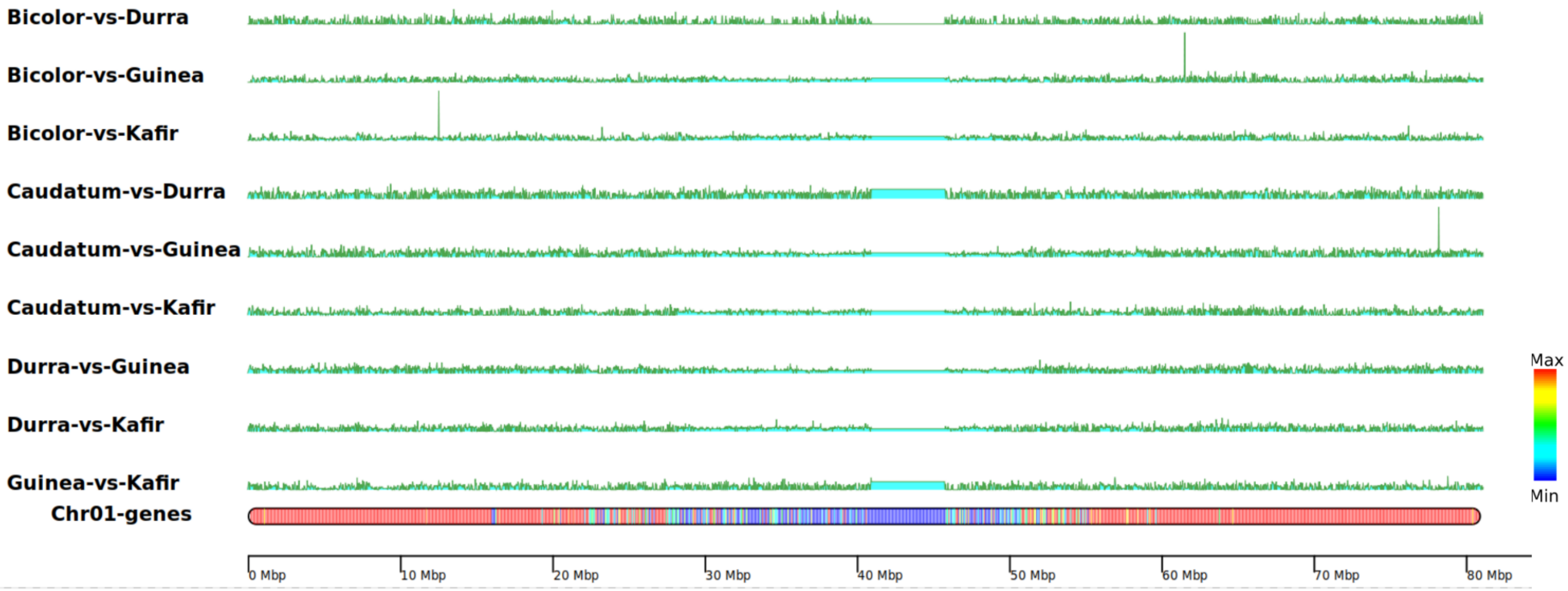
Bicolor-vs-Durra
Bicolor-vs-Guinea
Bicolor-vs-Kafir
Caudatum-vs-Durra
Caudatum-vs-Guinea
Caudatum-vs-Kafir
Durra-vs-Guinea
Durra-vs-Kafir
Guinea-vs-Kafir
Chr01-genes
0 Mbp
10 Mbp
20 Mbp
30 Mbp
40 Mbp
50 Mbp
60 Mbp
70 Mbp
80 Mbp
Max
Min

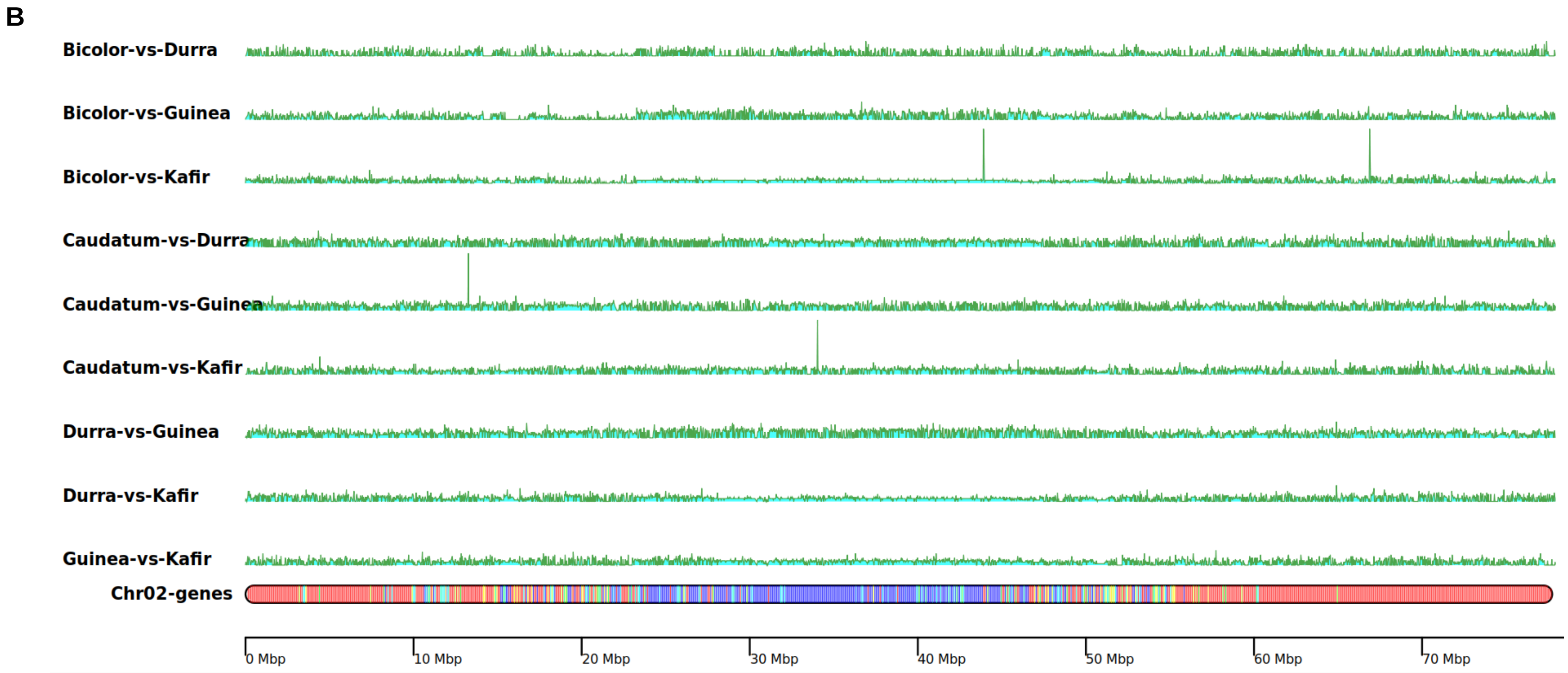
B
Bicolor-vs-Durra
Bicolor-vs-Guinea
Bicolor-vs-Kafir
Caudatum-vs-Durra
Caudatum-vs-Guinea
Caudatum-vs-Kafir
Durra-vs-Guinea
Durra-vs-Kafir
Guinea-vs-Kafir
Chr02-genes
0 Mbp
10 Mbp
20 Mbp
30 Mbp
40 Mbp
50 Mbp
60 Mbp
70 Mbp

C

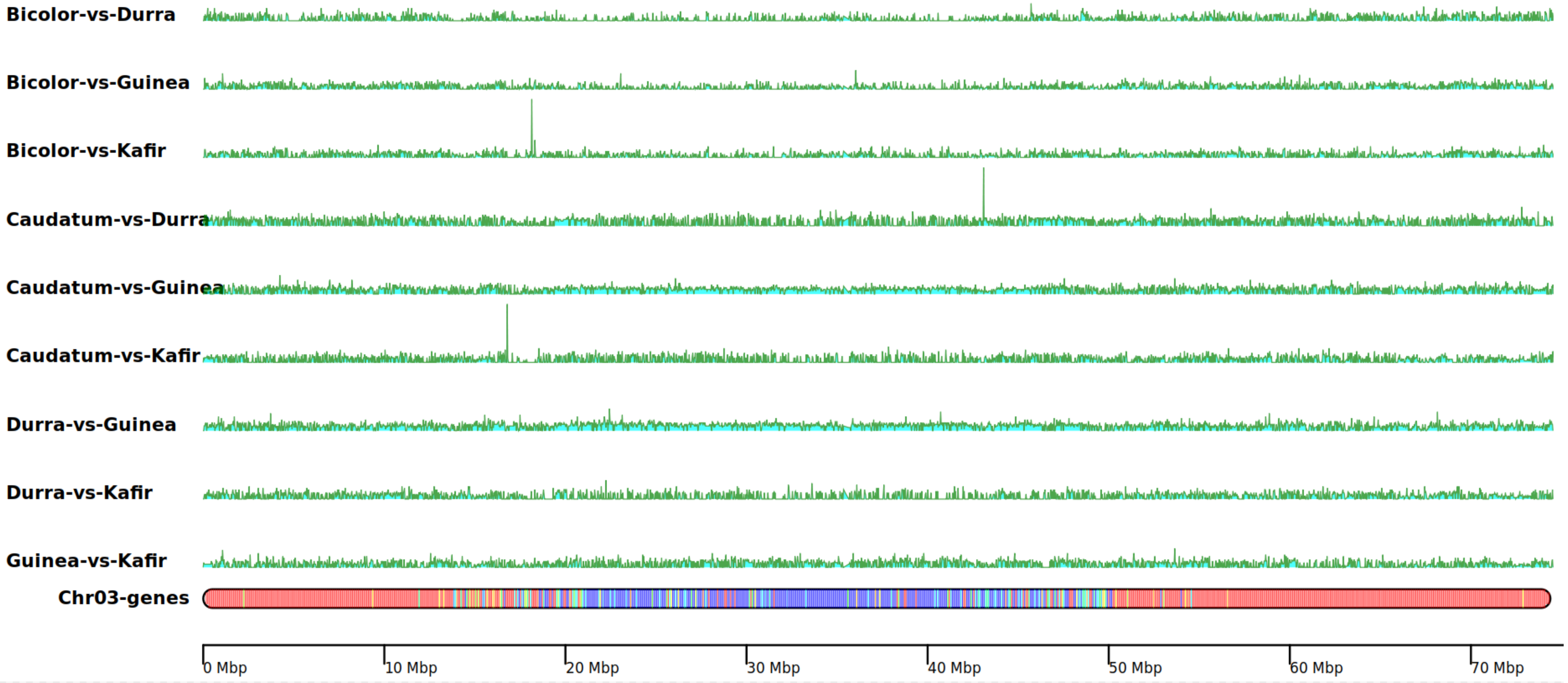

Bicolor-vs-Durra
Bicolor-vs-Guinea
Bicolor-vs-Kafir
Caudatum-vs-Durra
Caudatum-vs-Guinea
Caudatum-vs-Kafir
Durra-vs-Guinea
Durra-vs-Kafir
Guinea-vs-Kafir
Chr03-genes
0 Mbp
10 Mbp
20 Mbp
30 Mbp
40 Mbp
50 Mbp
60 Mbp
70 Mbp

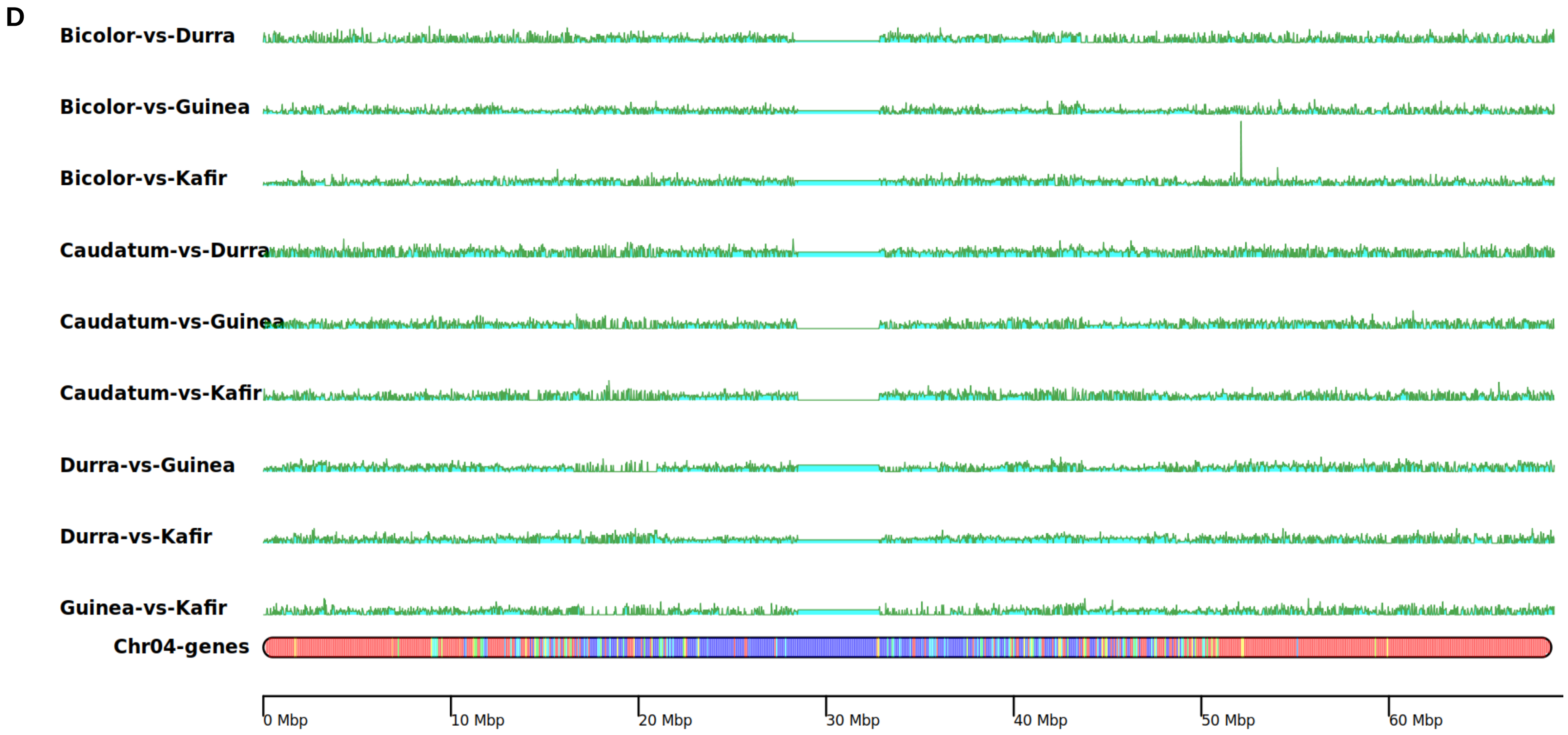
D
Bicolor-vs-Durra
Bicolor-vs-Guinea
Bicolor-vs-Kafir
Caudatum-vs-Durra
Caudatum-vs-Guinea
Caudatum-vs-Kafir
Durra-vs-Guinea
Durra-vs-Kafir
Guinea-vs-Kafir
Chr04-genes
0 Mbp
10 Mbp
20 Mbp
30 Mbp
40 Mbp
50 Mbp
60 Mbp

E

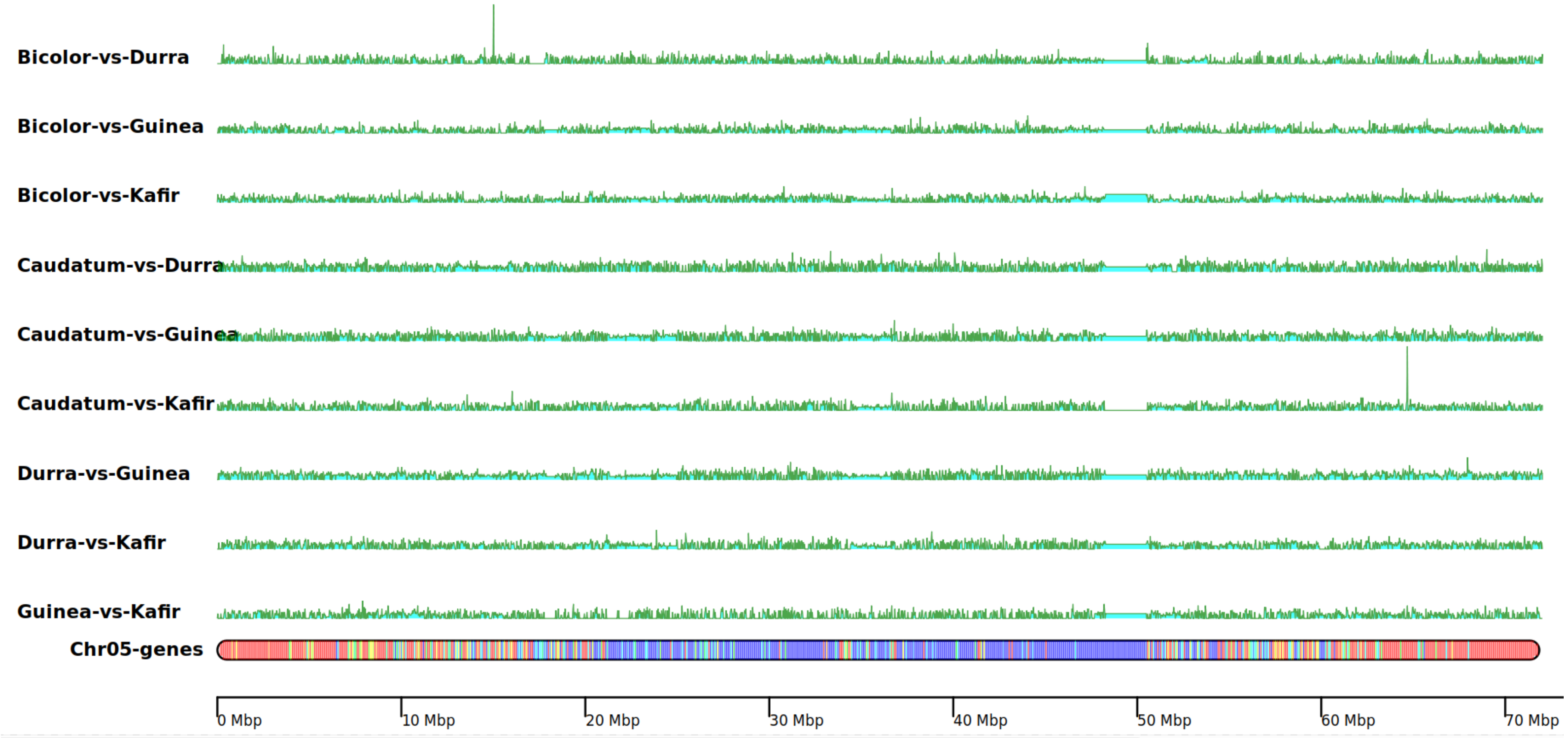

Bicolor-vs-Durra
Bicolor-vs-Guinea
Bicolor-vs-Kafir
Caudatum-vs-Durra
Caudatum-vs-Guinea
Caudatum-vs-Kafir
Durra-vs-Guinea
Durra-vs-Kafir
Guinea-vs-Kafir
Chr05-genes
0 Mbp
10 Mbp
20 Mbp
30 Mbp
40 Mbp
50 Mbp
60 Mbp
70 Mbp

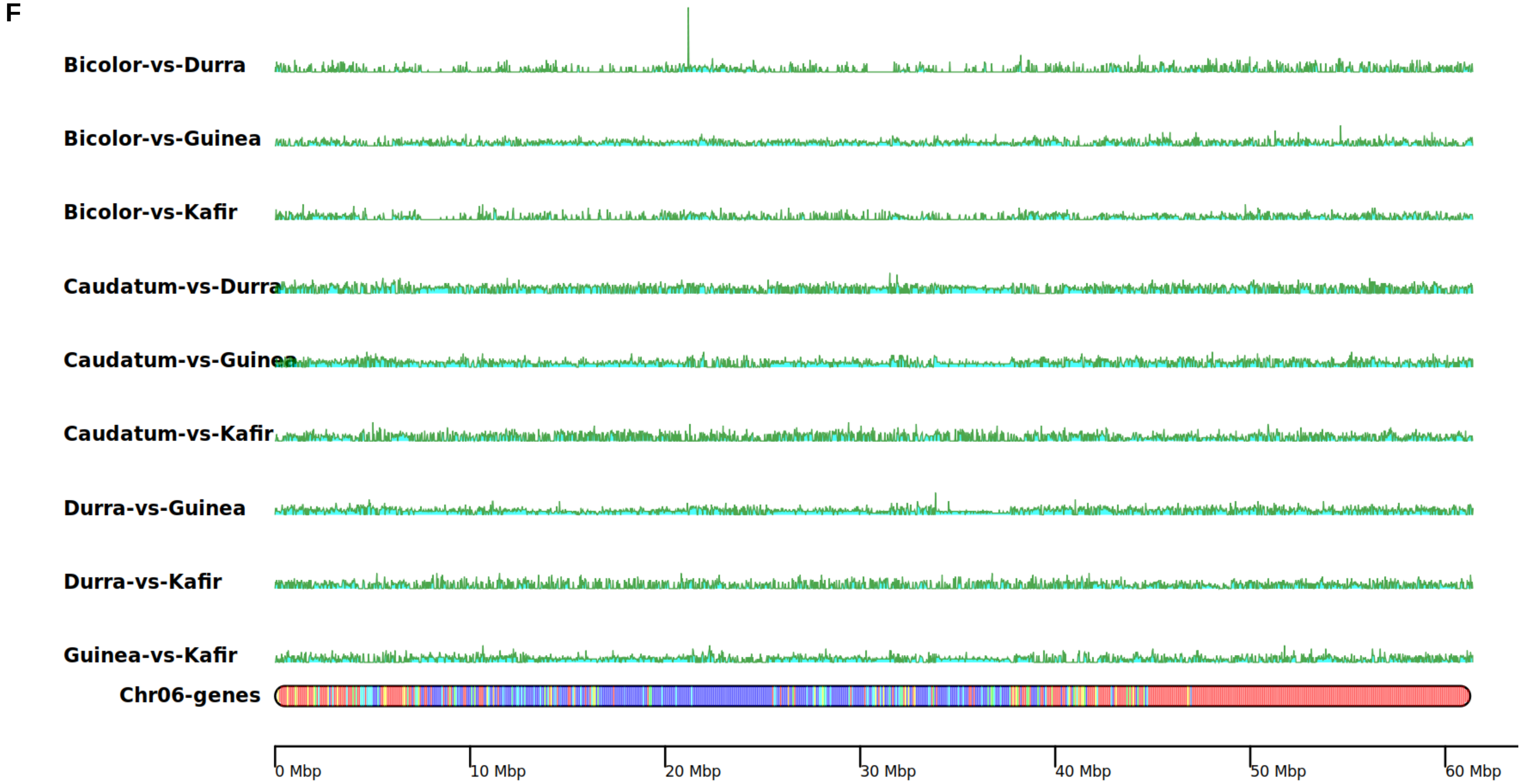
F
Bicolor-vs-Durra
Bicolor-vs-Guinea
Bicolor-vs-Kafir
Caudatum-vs-Durra
Caudatum-vs-Guinea
Caudatum-vs-Kafir
Durra-vs-Guinea
Durra-vs-Kafir
Guinea-vs-Kafir
Chr06-genes
0 Mbp
10 Mbp
20 Mbp
30 Mbp
40 Mbp
50 Mbp
60 Mbp

G

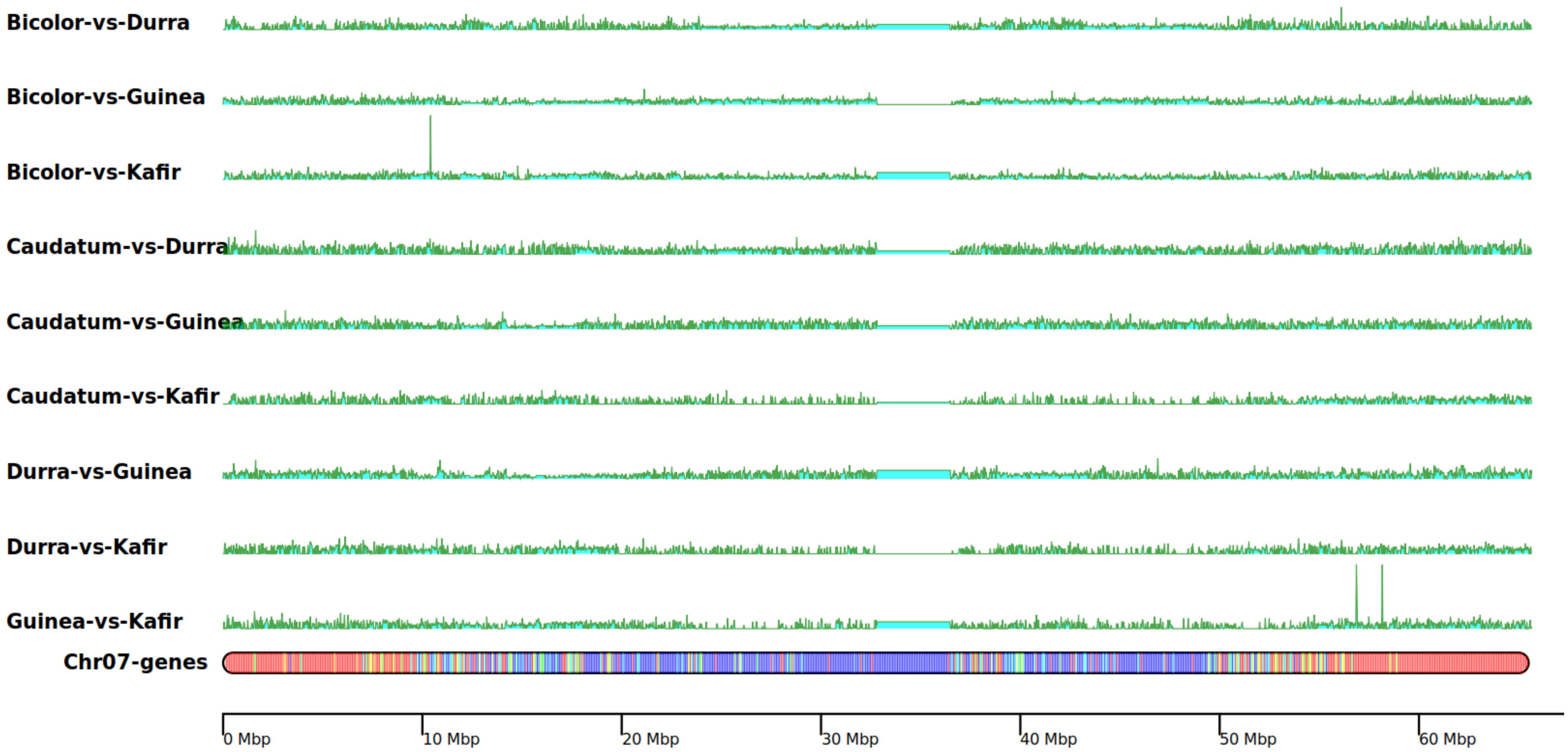
Bicolor-vs-Durra
Bicolor-vs-Guinea
Bicolor-vs-Kafir
Caudatum-vs-Durra
Caudatum-vs-Guinea
Caudatum-vs-Kafir
Durra-vs-Guinea
Durra-vs-Kafir
Guinea-vs-Kafir
Chr07-genes
0 Mbp
10 Mbp
20 Mbp
30 Mbp
40 Mbp
50 Mbp
60 Mbp

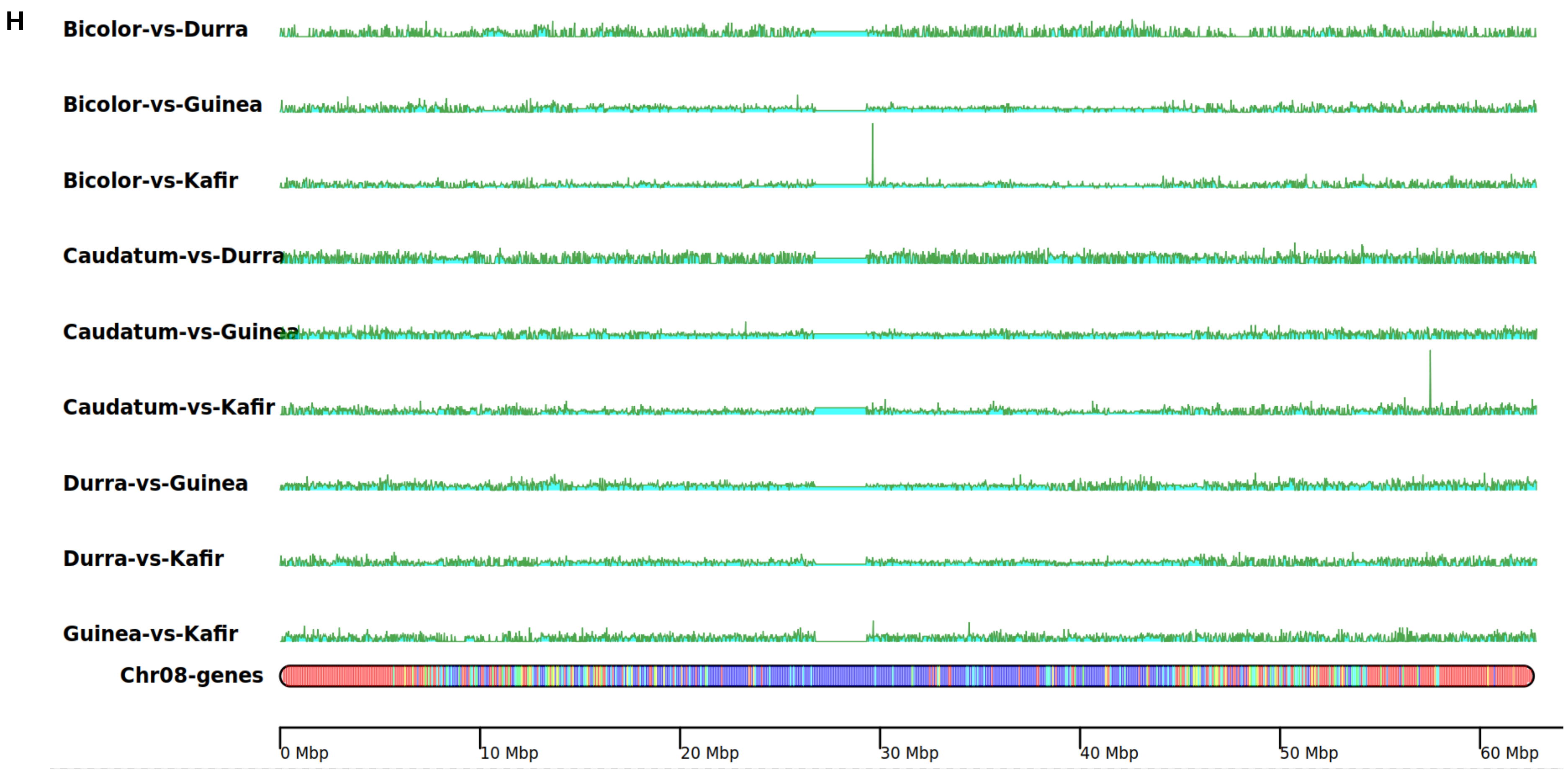
H
Bicolor-vs-Durra
Bicolor-vs-Guinea
Bicolor-vs-Kafir
Caudatum-vs-Durra
Caudatum-vs-Guinea
Caudatum-vs-Kafir
Durra-vs-Guinea
Durra-vs-Kafir
Guinea-vs-Kafir
Chr08-genes
0 Mbp
10 Mbp
20 Mbp
30 Mbp
40 Mbp
50 Mbp
60 Mbp

I

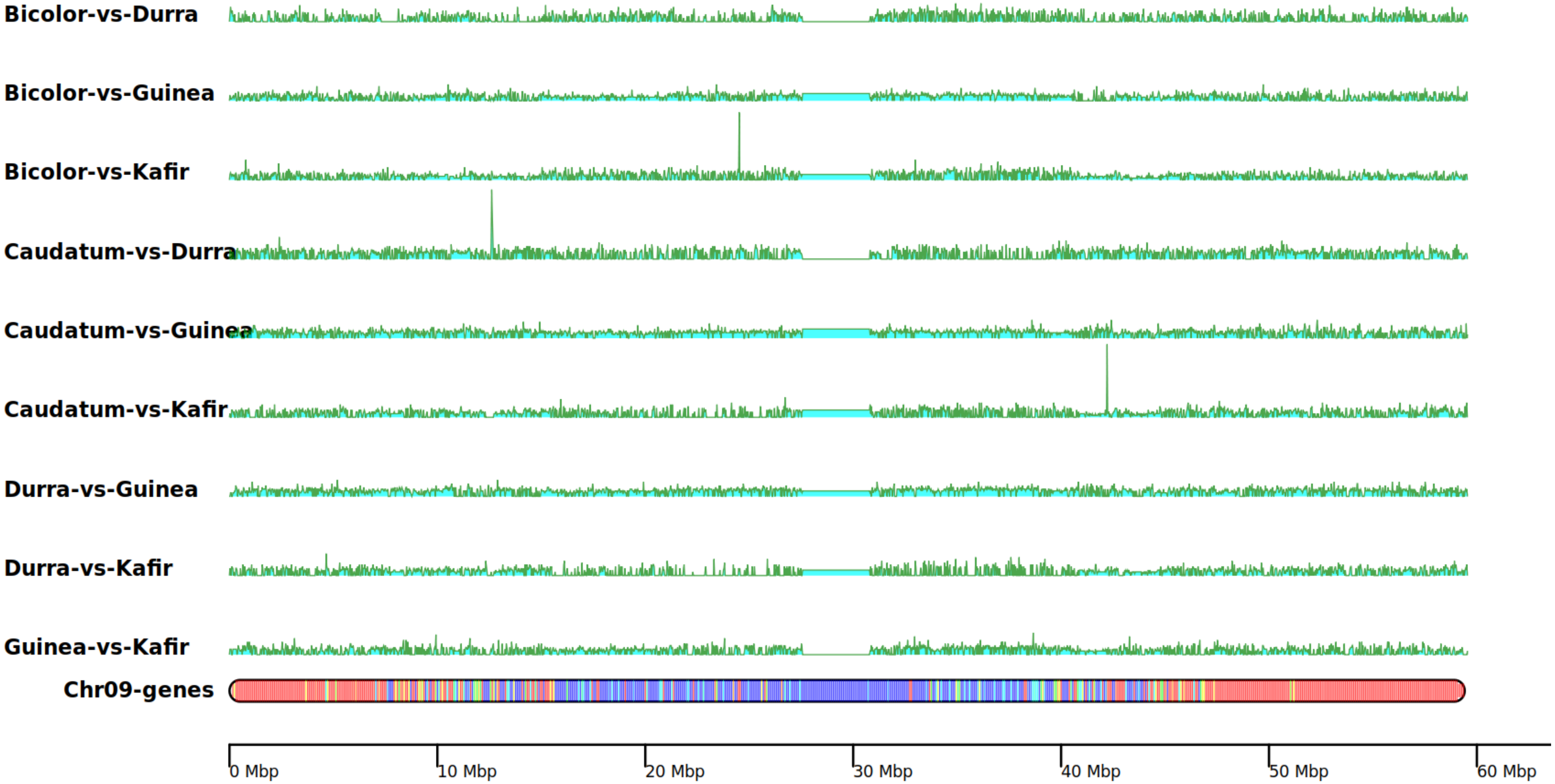
Bicolor-vs-Durra
Bicolor-vs-Guinea
Bicolor-vs-Kafir
Caudatum-vs-Durra
Caudatum-vs-Guinea
Caudatum-vs-Kafir
Durra-vs-Guinea
Durra-vs-Kafir
Guinea-vs-Kafir
Chr09-genes
0 Mbp
10 Mbp
20 Mbp
30 Mbp
40 Mbp
50 Mbp
60 Mbp

J

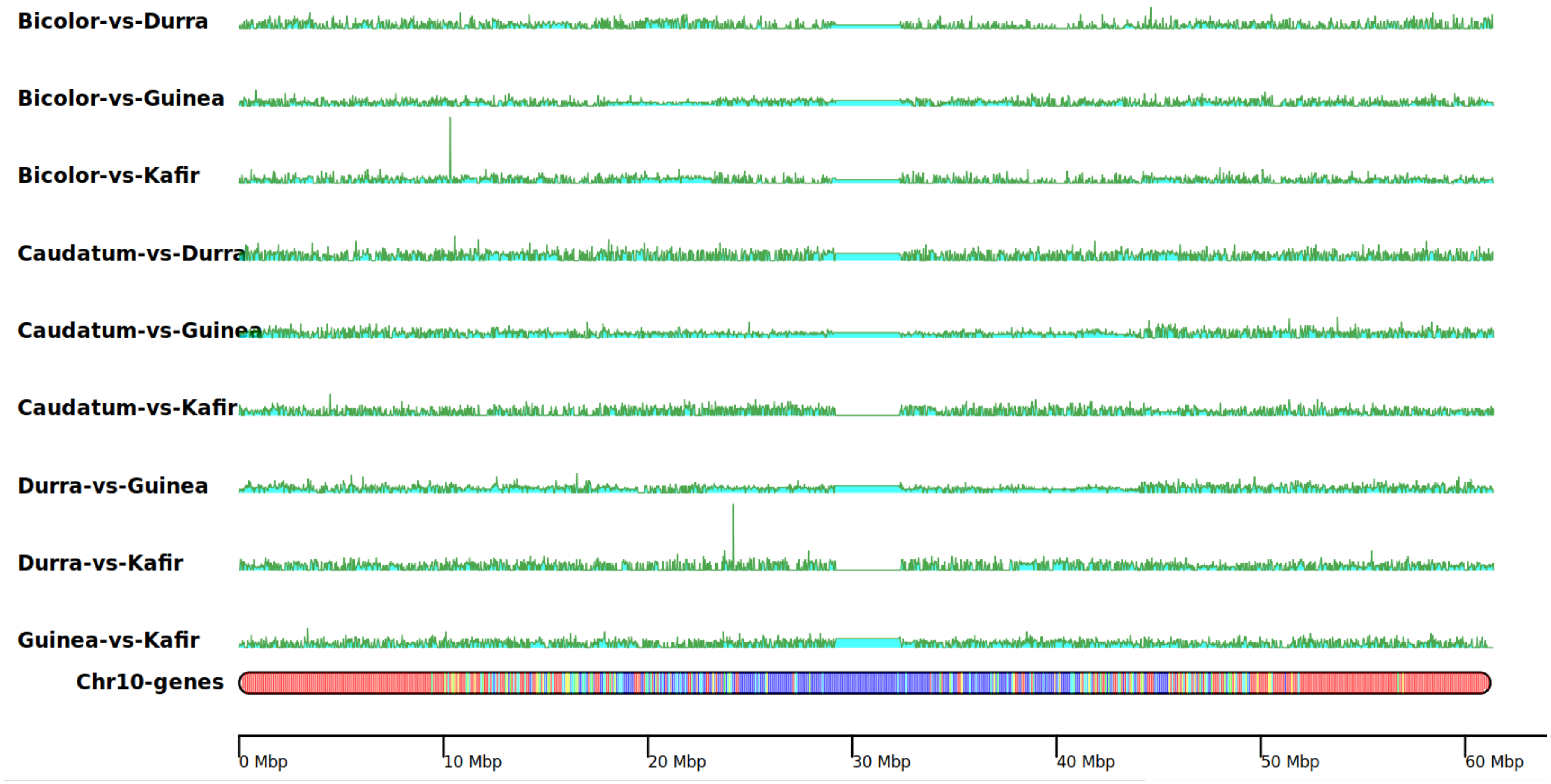


**Supplementary Figure 4.** Pairwise $F_{ST}$ across the chromosomes of sorghum race populations and gene density heatmap on respective chromosomes.

A

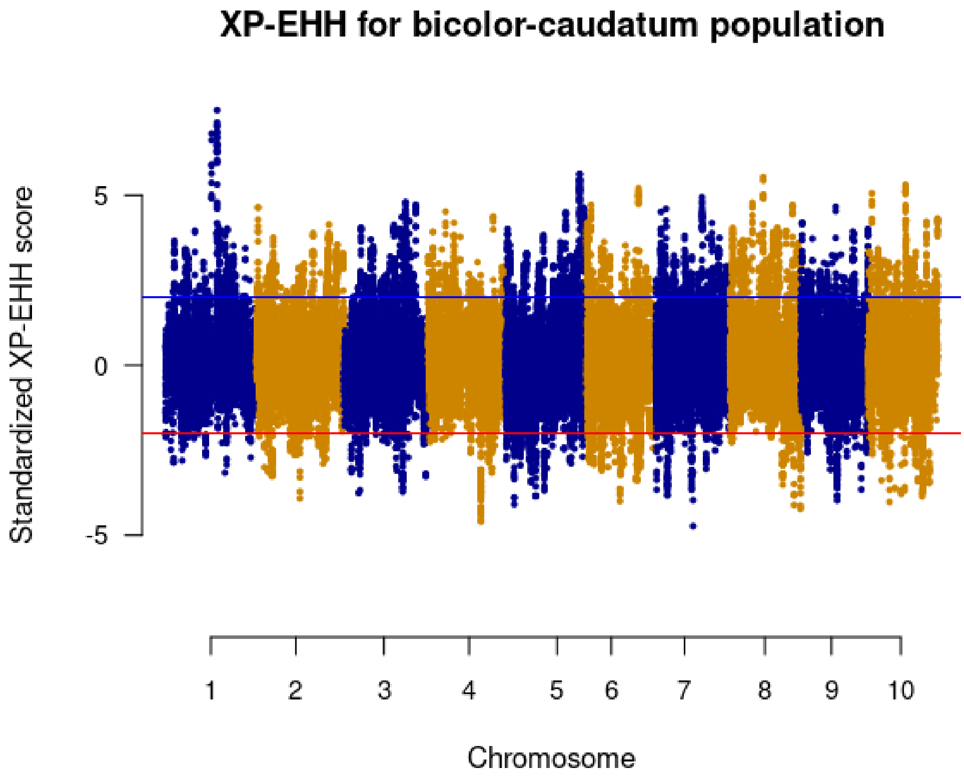
XP-EHH for bicolor-caudatum population
Standardized XP-EHH score
5
0
-5
1
2
3
4
5
6
7
8
9
10
Chromosome

B

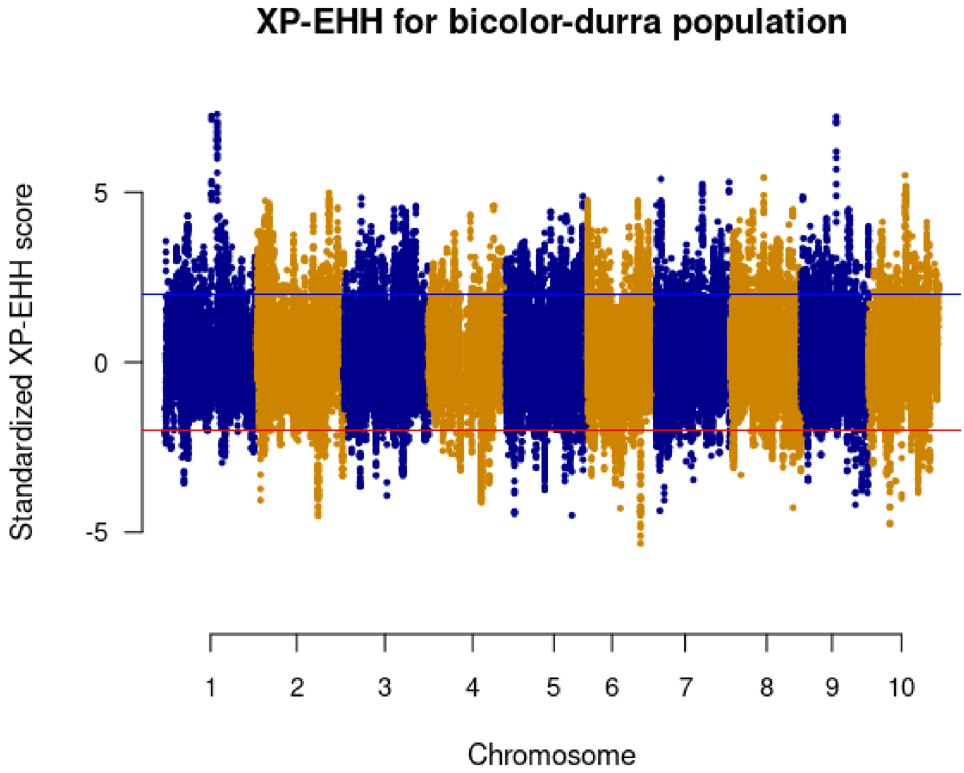
XP-EHH for bicolor-durra population
Standardized XP-EHH score
5
0
-5
1
2
3
4
5
6
7
8
9
10
Chromosome

C

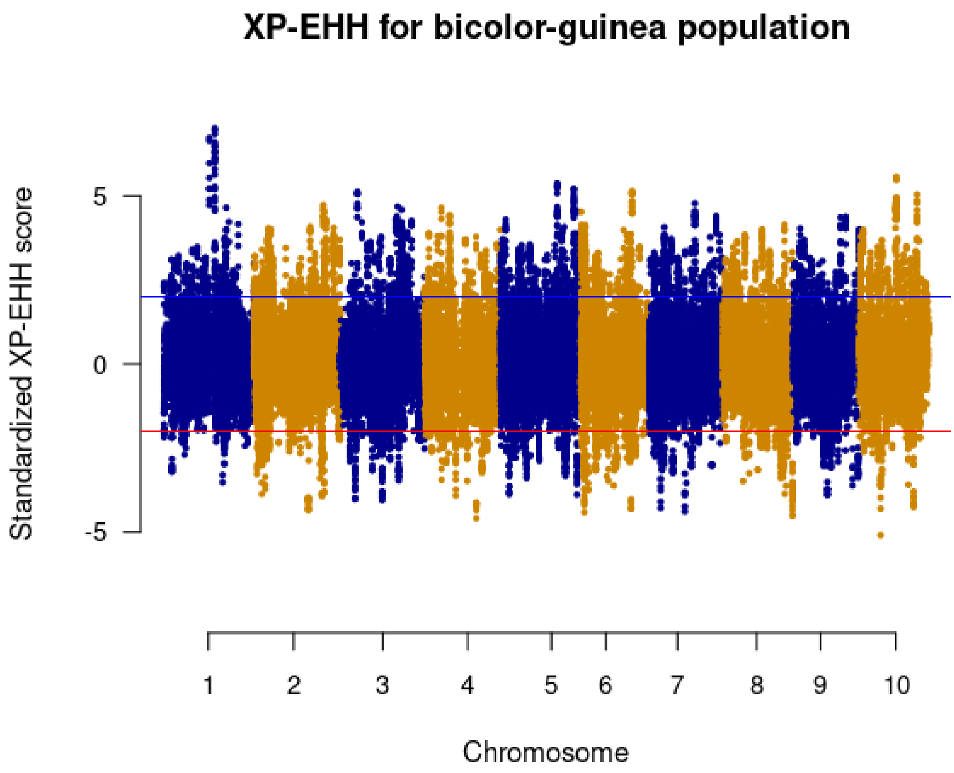
XP-EHH for bicolor-guinea population
Standardized XP-EHH score
5
0
-5
1
2
3
4
5
6
7
8
9
10
Chromosome

D

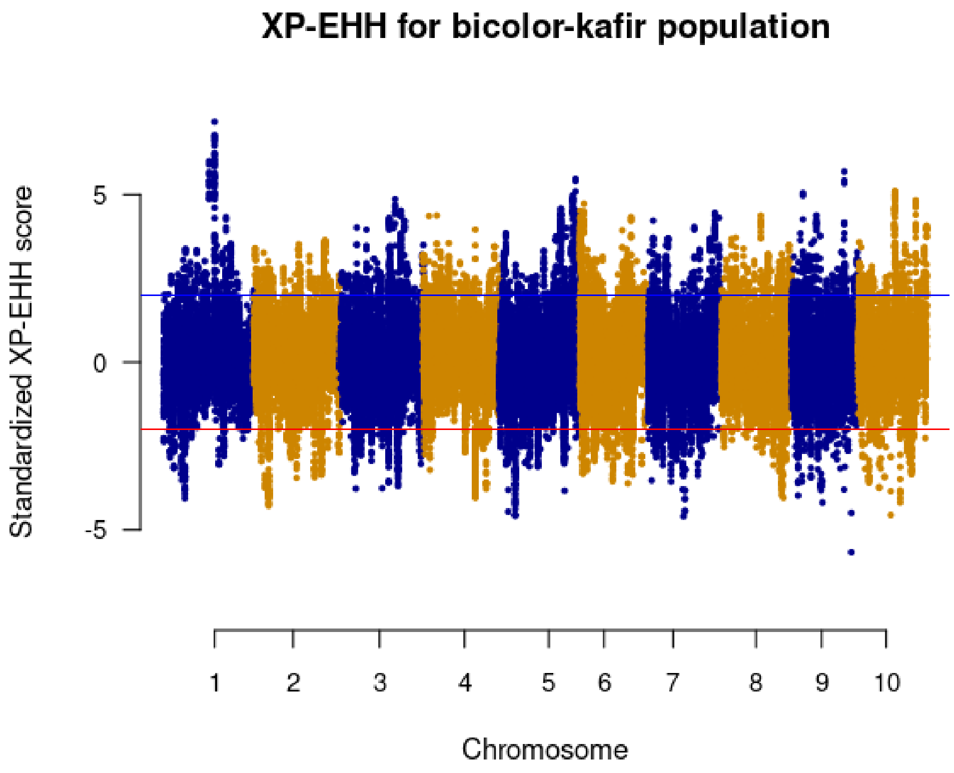
XP-EHH for bicolor-kafir population
Standardized XP-EHH score
5
0
-5
1
2
3
4
5
6
7
8
9
10
Chromosome

E

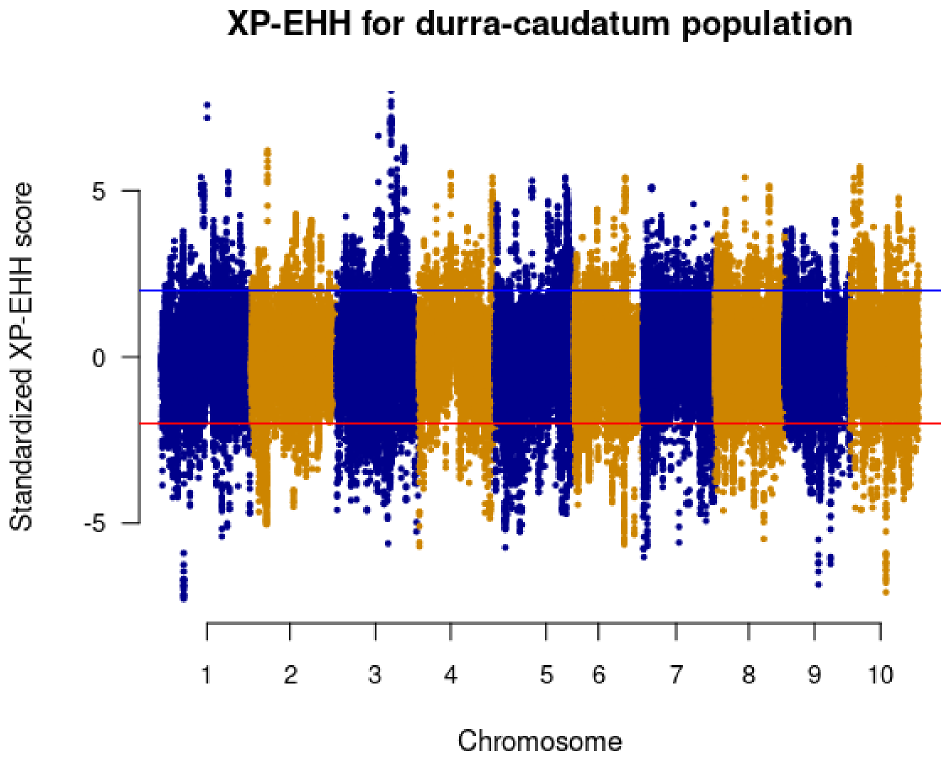
XP-EHH for durra-caudatum population
Standardized XP-EHH score
5
0
-5
1
2
3
4
5
6
7
8
9
10
Chromosome

F

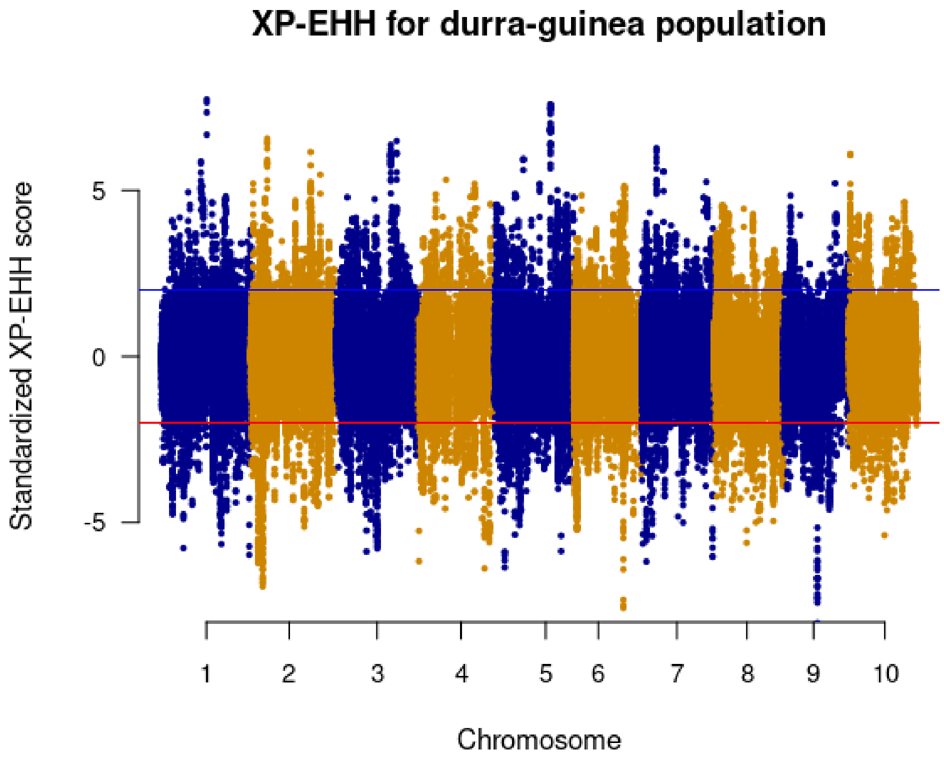
XP-EHH for durra-guinea population
Standardized XP-EHH score
5
0
-5
1
2
3
4
5
6
7
8
9
10
Chromosome

G

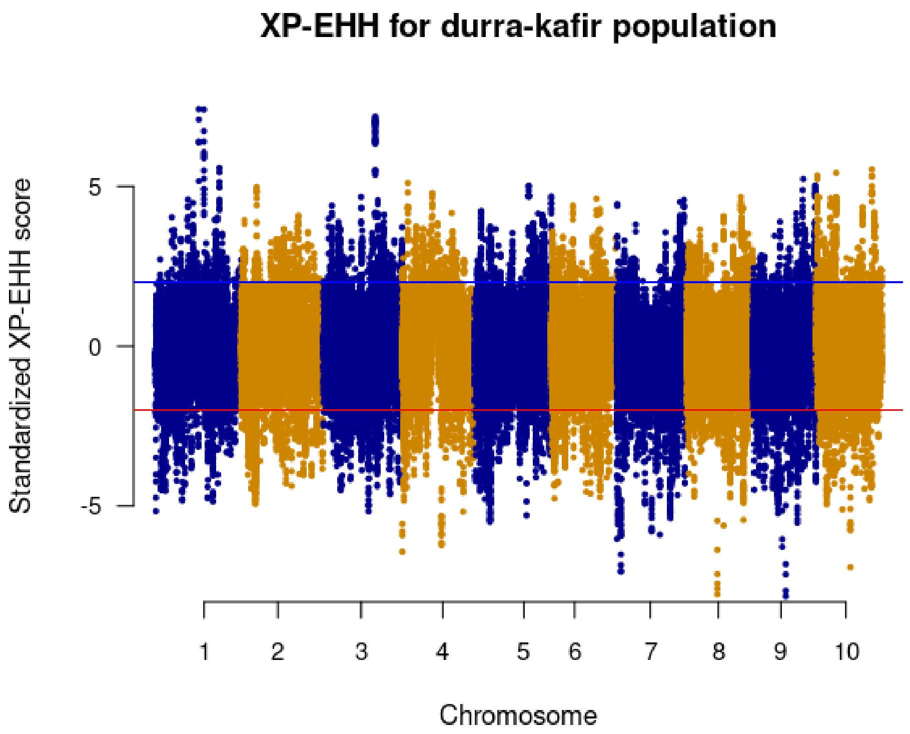
XP-EHH for durra-kafir population
Standardized XP-EHH score
5
0
-5
1
2
3
4
5
6
7
8
9
10
Chromosome

H

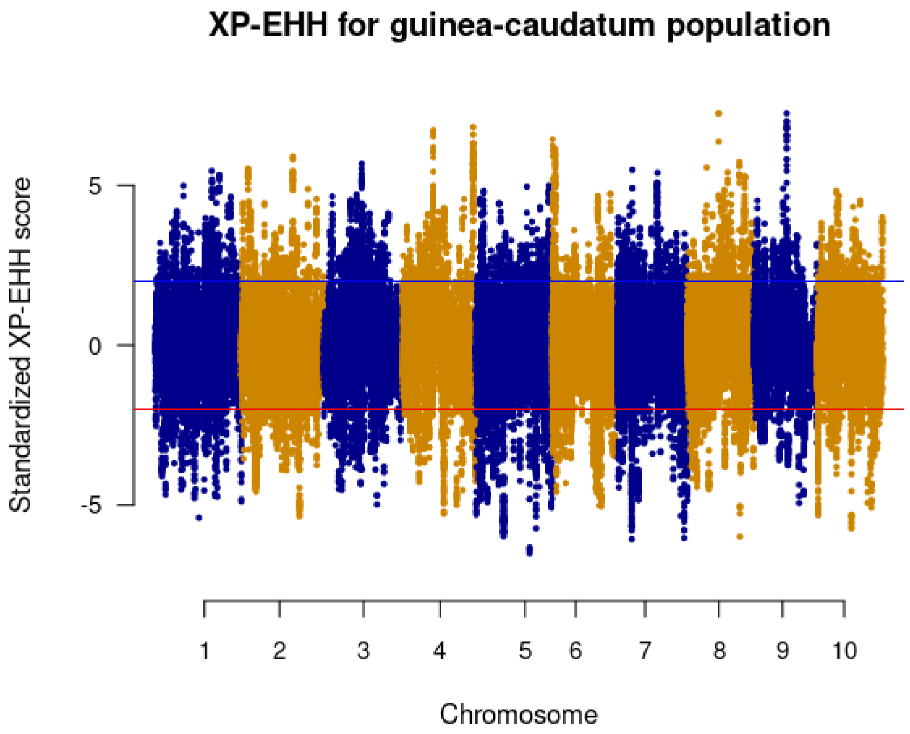
XP-EHH for guinea-caudatum population
Standardized XP-EHH score
5
0
-5
1
2
3
4
5
6
7
8
9
10
Chromosome

I

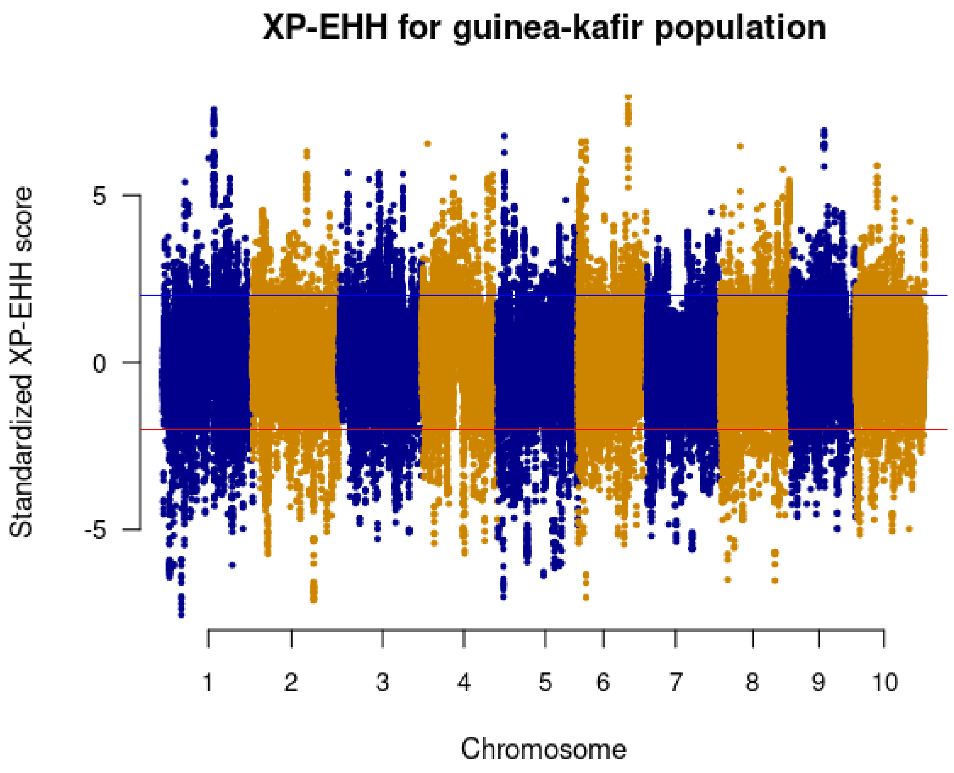


J

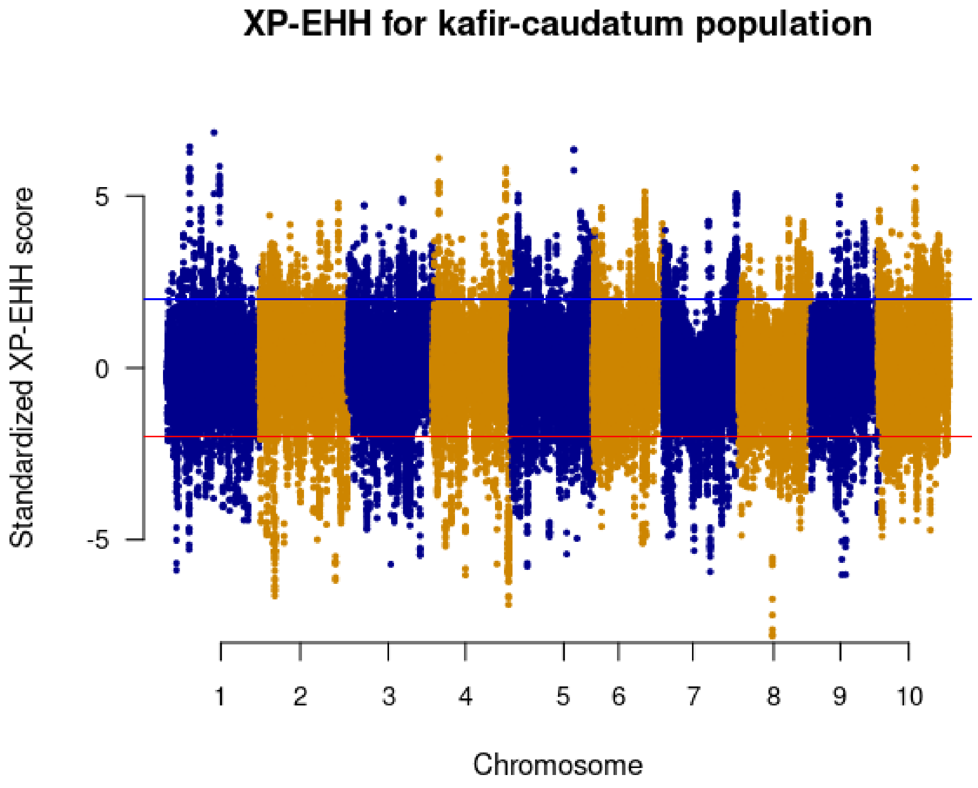


**Supplementary Figure 5.** Distribution of standardized XP-EHH scores in the sorghum race population comparison

A

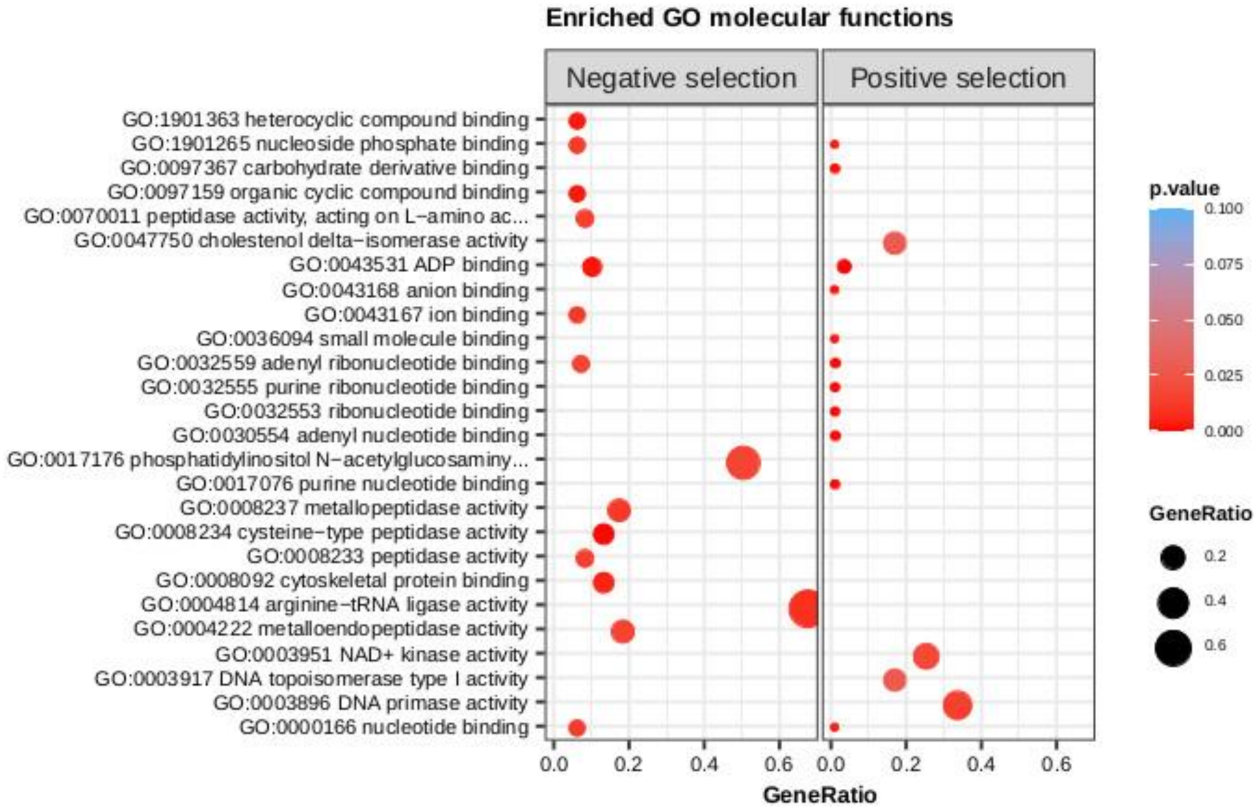

Enriched GO molecular functions
Negative selection
Positive selection
GO:1901363 heterocyclic compound binding
GO:1901265 nucleoside phosphate binding
GO:0097367 carbohydrate derivative binding
GO:0097159 organic cyclic compound binding
GO:0070011 peptidase activity, acting on L−amino ac...
GO:0047750 cholestenol delta−isomerase activity
GO:0043531 ADP binding
GO:0043168 anion binding
GO:0043167 ion binding
GO:0036094 small molecule binding
GO:0032559 adenyl ribonucleotide binding
GO:0032555 purine ribonucleotide binding
GO:0032553 ribonucleotide binding
GO:0030554 adenyl nucleotide binding
GO:0017176 phosphatidylinositol N−acetylglucosaminy...
GO:0017076 purine nucleotide binding
GO:0008237 metallopeptidase activity
GO:0008234 cysteine−type peptidase activity
GO:0008233 peptidase activity
GO:0008092 cytoskeletal protein binding
GO:0004814 arginine−tRNA ligase activity
GO:0004222 metalloendopeptidase activity
GO:0003951 NAD+ kinase activity
GO:0003917 DNA topoisomerase type I activity
GO:0003896 DNA primase activity
GO:0000166 nucleotide binding
0.0
0.2
0.4
0.6
GeneRatio
p.value
0.100
0.075
0.050
0.025
0.000
GeneRatio
0.2
0.4
0.6

B

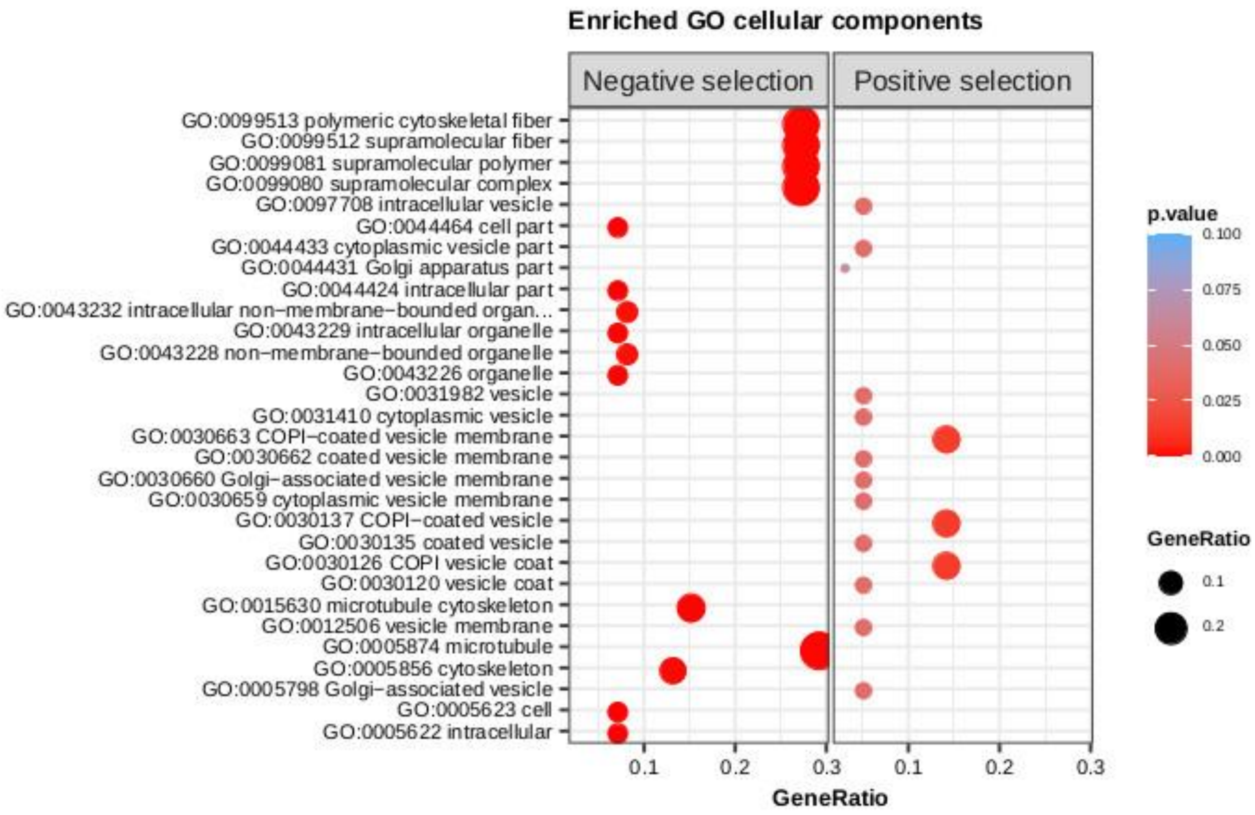


**Supplementary Figure 6.** Enrichment of genes under selection pressure for **A)** molecular function **B)** Cellular components.
